# What is an adaptive pattern of brain network coupling for a child? It depends on their environment

**DOI:** 10.1101/2020.05.29.124297

**Authors:** Monica E. Ellwood-Lowe, Susan Whitfield-Gabrieli, Silvia A. Bunge

## Abstract

Prior research indicates that lower resting-state functional coupling between two brain networks, lateral frontoparietal network (LFPN) and default mode network (DMN), relates to better cognitive test performance. However, most study samples skew towards wealthier individuals—and what is adaptive for one population may not be for another. In a pre-registered study, we analyzed resting-state fMRI from 6839 children ages 9-10 years. For children above poverty, we replicated the prior finding: better cognitive performance correlated with weaker LFPN-DMN coupling. For children in poverty, the slope of the relation was instead positive. This significant interaction related to several features of a child’s environment. Future research should investigate the possibility that leveraging internally guided cognition is a mechanism of resilience for children in poverty. In sum, “optimal” brain function depends in part on the external pressures children face, highlighting the need for more diverse samples in research on the human brain and behavior.

## Introduction

In the United States, one fifth of children are estimated to live below the poverty line (Semega et al., 2019). Relative to children living just above poverty, these children are least likely to have access to the federal social safety net, and they are at heightened risk for poor health and educational outcomes (Hoynes & Schanzenbach, 2018; Reardon, 2016). Compared to their peers whose families earn more money, children living in poverty tend to perform worse on tests of cognitive functioning (for a review, see Farah, 2017), itself a risk factor for later outcomes (e.g., Spengler et al., 2015). However, such broad comparisons obscure substantial variability *within* the group of children living in poverty, a large segment of whom score on par with their higher-income peers. Here, we seek to understand this form of resilience—high cognitive test performance in the face of structural barriers to success. One way to begin to address this question is to identify sets of experiences that may be protective for children in poverty, given the wide range of experiences they have (DeJoseph et al., 2020; Gonzalez et al., 2019). Another way is to probe differences in brain function, to gain insight into the mechanisms underlying resilience. In this study, we examine the neural and environmental correlates of resilience in a sample of over 1,000 children across the United States likely to be living in poverty.

In one of the most influential theories of development, Waddington proposed that ontogenetic trajectories are variable across individuals and not inherently fixed at birth (Johnson & de Haan, 2015; Waddington, 1957). Instead, both biological and environmental influences interact across development to constrain the ultimate expression of cells in our bodies. This means that in some cases, environmental pressures, especially early in life, may cause two individuals with the same biological constraints to develop different phenotypes. In other cases, two individuals may take distinct developmental trajectories, but ultimately still develop the same phenotype (Edelman & Gally, 2001). Extending this metaphor to the current study, it is possible that two children who display the same level of performance on a cognitive test might achieve this through different developmental trajectories, if they grow up under different external pressures. The optimal developmental trajectory for a child, therefore, may be influenced by the child’s environment.

Accumulating evidence suggests that the brain adapts to the affordances and constraints of an individual’s environment, especially in early life. Indeed, a growing number of studies have complicated the notion of an “ideal” environment by suggesting that different environments promote the development of distinct, adaptive cognitive skills (Frankenhuis et al., 2019; Mittal et al., 2015; Young et al., 2018) The result of this adaptability may be that higher-level cognitive skills such as executive functions and reasoning, which build on lower-level skills that may be more environmentally sensitive, develop in context-sensitive ways. Children living in poverty can have vastly different experiences than those who are typically studied in developmental cognitive neuroscience, including varying levels of threat exposure and resource deprivation (Humphreys & Zeanah, 2015; McLaughlin et al., 2014). Understanding the ways in which their brains may have been tuned by their respective environments can provide insight into mechanisms of adaptation, and, ultimately, how best to support each child within the specific constraints of their lives.

Strikingly, while much research has characterized the trajectories of brain development that support cognitive test performance for upper-middle class children— most of whom who tend to be living in urban places close to universities in the United States—only in the last decade has research begun to focus on children from lower socioeconomic status (SES) backgrounds. This new thrust of research has begun to uncover neural differences between higher- and lower-SES children in brain structure and function from an early age (e.g., Hair et al., 2015; Hanson et al., 2013; S. B. Johnson et al., 2016; Leonard et al., 2019; Mackey et al., 2015; Noble et al., 2015; Noble et al., 2006). However, even in this literature, children living below the poverty line tend to be under-represented. In addition, many studies compare higher and lower SES children, obscuring variability within the lower SES group. Thus, characterizing optimal brain development for children living below poverty could help shift our questions away from how these children differ from children above poverty, and toward understanding mechanisms supporting neurocognitive functioning in an understudied population. Ultimately, this brings us toward a fuller understanding of brain development across the full spectrum of life experiences.

In line with the hypothesis that children may achieve the same behavior or phenotype through different developmental routes, studies examining brain function during higher-level cognitive tasks often find qualitatively different brain-behavior relations as a function of children’s family income. Differences in brain activation appear particularly in lateral prefrontal cortex (PFC) and parietal regions—two regions that are involved in higher cognitive function, show protracted development (Casey et al., 2000), and are sensitive to environmental input (Farah, 2017; Mackey et al., 2013; Merz, Maskus, et al., 2019).

Collectively, these and other studies suggest that children with lower versus higher family incomes may differentially engage higher-order brain areas such as lateral prefrontal and parietal regions to complete tasks that tax working memory, rule learning, and attention (Finn et al., 2017; Sheridan et al., 2012; see Merz, Wiltshire, & Noble, 2019 for a review). These differences in brain function are typically thought to reflect differences in either the cognitive mechanisms by which children approach the task or efficiency of neural processing. However, differences in tasks and task demands make it difficult to generalize across studies showing divergent prefrontal and parietal activation as a function of SES. Interpretation of differences in brain function during performance of a specific task is constrained by task demands. For example, there may be unseen verbal demands that differentially affect some children’s approach to the task more than others’; additionally, the tasks are not representative of real-world experiences, limiting validity.

Another way to investigate SES differences in brain function is to measure slow-wave fluctuations in neural activity over time while participants lie awake in an MRI scanner, in the absence of specific task demands. This approach, called resting-state fMRI, has revealed temporal coupling among anatomically distal brain regions that form large-scale brain networks (Uddin et al., 2019). In general, cognitive networks become more cohesive and segregated from one another across development (Grayson & Fair, 2017; Power et al., 2010). Patterns of temporal coupling within and across resting-state networks reflect regions’ prior history of co-activation, offering insight into individuals’ recent thought pattern (Guerra-Carrillo et al., 2014). Thus, resting-state fMRI can be leveraged to assess how everyday experience shapes brain networks. With regard to SES, there is evidence that children and adolescents living in disadvantaged neighborhoods show differences in resting-state connectivity patterns, some of which correlate with anxiety symptomatology (Marshall et al., 2018). Further, changes in family income in adolescence have been associated with changes in connectivity in frontal and parietal regions associated primarily with the default mode network (Weissman et al., 2018). It is important to understand both how these differences arise and the ways in which they are behaviorally relevant.

Several large-scale brain networks have been linked to higher-level cognition (Barber et al., 2013; Hampson et al., 2010; Keller et al., 2015; Kelly et al., 2008). In particular, the lateral frontoparietal network (LFPN) is consistently activated in higher-level cognitive tasks, such as those taxing executive functions or reasoning. Regions in the LFPN are more active during performance of cognitively demanding tasks than during rest periods (Vincent et al., 2008). In contrast, regions in the default mode network (DMN), including regions in the medial frontal and medial parietal areas, are consistently de-activated during focused task performance. These regions have been implicated in unconstrained, internally directed thought (Raichle et al., 2001), as well as during performance of tasks that require introspection, mentalizing about others, or other mentation outside of the here-and-now (Spreng, 2012). In fact, elevated DMN activation during performance of tasks that require focused attention has been associated with lower task accuracy and response times, and higher response variability (Kelly et al., 2008; Satterthwaite et al., 2013; D. H. Weissman et al., 2006).

Thus, the LFPN and DMN have often been characterized as opponent networks. Indeed, a number of studies of young adults have linked weaker resting-state connectivity between the LFPN and DMN, and stronger connectivity among LFPN regions, to better cognitive performance (Barber et al., 2013; Hampson et al., 2010; Keller et al., 2015; Kelly et al., 2008). These findings suggest that, in order to complete a cognitively demanding task, individuals must focus narrowly on the task at hand while inhibiting internally-directed or self-referential thoughts (Raichle et al., 2001; Simpson et al., 2001a, 2001b; D. H. Weissman et al., 2006).

This conclusion has been bolstered by fMRI research in typically developing children, both in terms of age-related changes and individual differences. First, there is evidence that the LFPN and DMN functionally segregate during childhood. Specifically, key nodes in the LFPN and DMN have been shown to be positively correlated in middle childhood, anti-correlated in adolescence, and more strongly anti-correlated during young adulthood (Chai et al., 2014b). Further, as with adults, children ages 10-13 who showed less coupling than their same-age peers tended to have higher cognitive task scores (Sherman et al., 2014). Tighter coupling between key nodes in these networks at age 7 has even been shown to predict increased attentional problems over the subsequent four years (Whitfield-Gabrieli et al., 2020). The conclusion drawn from these studies is that it is adaptive for LFPN and DMN to become decoupled—or even negatively coupled—during performance of a cognitively challenging task, and that the development of this dissociation may promote stronger focus on externally directed tasks.

Despite this coherent body of findings regarding these networks and their interactions, several points bear mentioning. First, there is evidence that LFPN and DMN interact during performance of tasks that benefit from internally directed cognition, or mentation outside of the here-and-now (Buckner & Carroll, 2007; Christoff et al., 2009; Kam et al., 2019; Spreng, 2012). Second, the vast majority of fMRI studies involve relatively high SES samples; thus, we do not know whether the reported brain-behavior relations are universal. Here, we sought to test the relation between connectivity of these two networks and cognitive task performance in a new sample: children living in poverty.

Drawing from a large behavioral and brain imaging dataset including over 10,000 children across the United States (ABCD Study; Casey et al., 2018), we asked whether the patterns of connectivity that are adaptive among higher-SES children also help to explain why some children living in poverty perform as well on cognitive tasks as their higher-income peers. Specifically, in a set of pre-registered analyses, we tested whether characteristics of LFPN and DMN connectivity were associated with cognitive test performance for over 1,000 children from this larger dataset who were estimated to be living in poverty. We sought to capture children’s performance on higher-level cognitive tasks that did not task verbal skills, given well-established SES differences in verbal performance. Thus, we combined measures of children’s abstract reasoning (Matrix reasoning task), inhibitory control (Flanker task), and cognitive flexibility (Dimensional Change Card Sort task).

Given prior evidence from higher-SES children and adults, we predicted that weaker LFPN-DMN between-network connectivity (decreased LFPN-DMN temporal coupling) and stronger within-network LFPN connectivity (LFPN-LFPN coupling) would be related to higher cognitive test performance even for children living in poverty. Alternatively, however, children in poverty might develop different brain-behavior links in order to contend with different barriers. In line with theories that children could achieve the same phenotype through alternate developmental trajectories, one might expect that higher cognitive test scores would be associated with different patterns of network connectivity among children in poverty. To preview our findings, our analyses revealed a different pattern in children in poverty than had been observed in prior studies of higher SES children. As a result, we conducted follow-up analyses involving the higher-income children in this sample to test whether their data would replicate prior findings, and confirmed that it did.

In a second set of pre-registered analyses, we probed demographic variables to better understand features of children’s environments which might explain variability both in their cognitive test performance, and in the relation between LFPN-DMN connectivity and cognitive test performance. We looked at a set of 29 variables that span home, school, and neighborhood contexts to see whether they could predict variability in children in poverty’s test performance. We also included interactions between LFPN-DMN connectivity and each of these variables, to see if patterns of brain-behavior relations could be explained by any particular set of experiences.

This study examines brain development in a large sample of children living below the poverty line. These children had a total family income below $35,000 (below $25,000 for children in families of 4 or less), a departure from the sample composition of most prior studies. Moreover, the tight age range in this dataset—all children were between 9 and 10 years old—complements prior studies of SES differences in brain development that have considered children across a much wider age range. Ultimately, examining relations between patterns of brain activity and cognitive test performance could help to elucidate the mechanisms through which high-performing children in poverty are able to contend with structural barriers in their environments.

## Results

We identified 1,034 children between ages 9 and 10 with usable data on cognitive test performance, resting state fMRI, and demographic characteristics, who were likely to be living below the poverty line at the time the data were collected (2016-2018). We identified an additional 5,805 children from the same study sites who had usable data on the same measures and were likely to be living *above* the poverty line. Participant information is displayed in Tables 1 and 2.

**Table 1.**
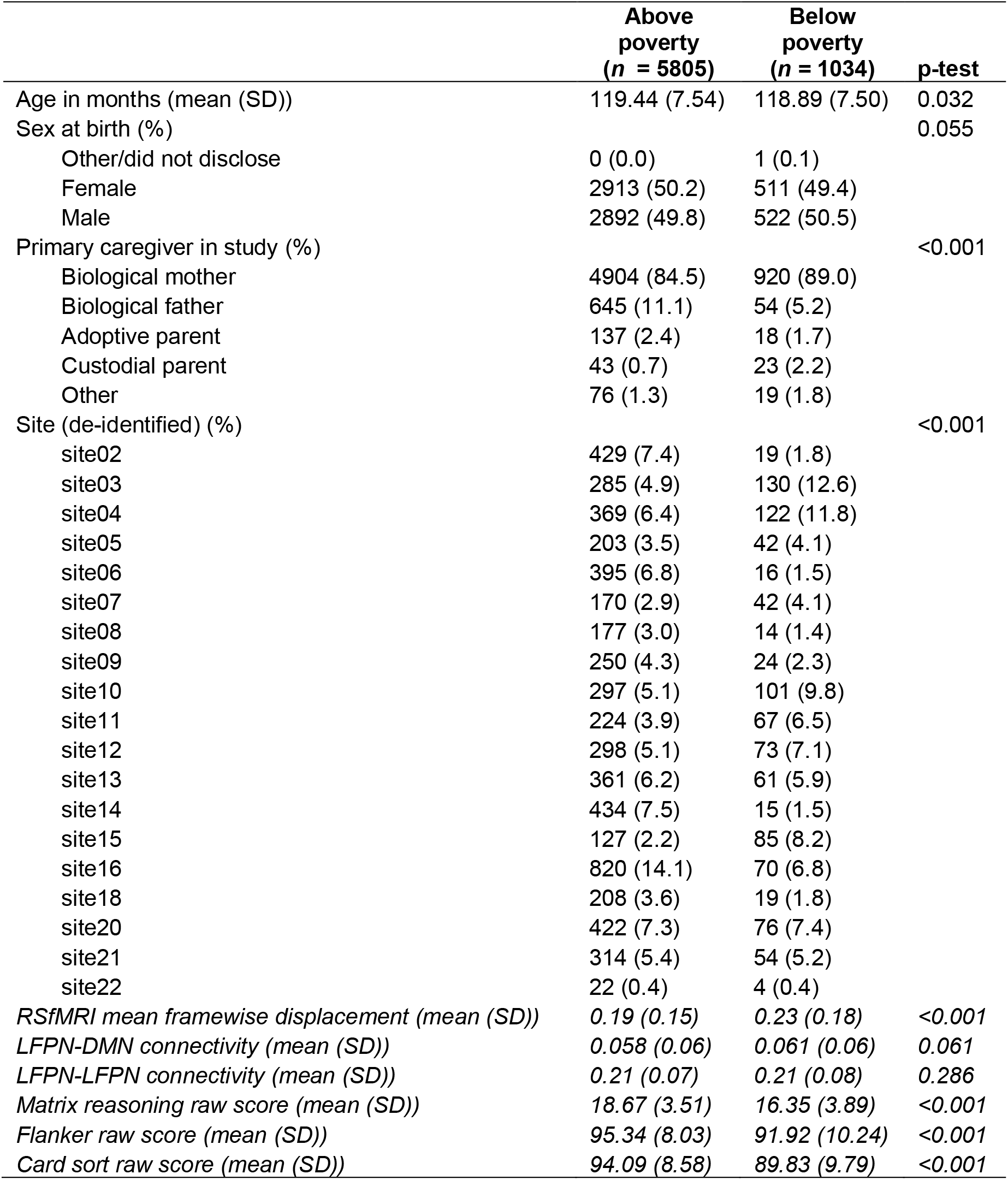
Participant characteristics. Demographic information in plain text; brain and cognitive variables italicized.

**Table 2.** Wider environmental information. Variables included in the ridge regression predicting cognitive test scores. All except income were used in primary models; additional tests confirmed that income did not add predictive power above and beyond these variables.

|  | Above poverty<br>(*n* = 5805) | Below poverty<br>(*n* = 1034) | p-test |
| --- | --- | --- | --- |
| Combined family income (%) |  |  | <0.001 |
| Less than \$5,000 | 0 (0.0) | 187 (18.1) | |
| \$5,000 through 11,999 | 0 (0.0) | 219 (21.2) | |
| \$12,000 through \$15,999 | 0 (0.0) | 154 (14.9) | |
| \$16,000 through \$24,999 | 0 (0.0) | 280 (27.1) | |
| \$25,000 through \$34,999 | 215 (3.7) | 194 (18.8) | |
| \$35,000 through \$49,999 | 579 (10.0) | 0 (0.0) | |
| \$50,000 through \$74,999 | 972 (16.7) | 0 (0.0) | |
| \$75,000 through \$99,999 | 1050 (18.1) | 0 (0.0) | |
| \$100,000 through \$199,999 | 2157 (37.2) | 0 (0.0) | |
| \$200,000 and greater | 832 (14.3) | 0 (0.0) | |
| Parents' highest level of education (n, %) |  |  | <0.001 |
| 3rd grade | 1 (0.0) | 0 (0.0) |  |
| 4th grade | 0 (0.0) | 1 (0.1) |  |
| 5th grade | 0 (0.0) | 1 (0.1) |  |
| 6th grade | 4 (0.1) | 13 (1.3) |  |
| 7th grade | 1 (0.0) | 2 (0.2) |  |
| 8th grade | 1 (0.0) | 8 (0.8) |  |
| 9th grade | 6 (0.1) | 24 (2.3) |  |
| 10th grade | 10 (0.2) | 26 (2.5) |  |
| 11th grade | 12 (0.2) | 34 (3.3) |  |
| 12th grade | 13 (0.2) | 47 (4.5) |  |
| High school graduate | 167 (2.9) | 169 (16.3) |  |
| GED or equivalent | 66 (1.1) | 91 (8.8) |  |
| Some college | 590 (10.2) | 297 (28.7) |  |
| Associate degree: occupational | 374 (6.4) | 135 (13.1) |  |
| Associate degree: academic | 297 (5.1) | 63 (6.1) |  |
| Bachelor's degree | 1818 (31.3) | 86 (8.3) |  |
| Master's degree | 1677 (28.9) | 32 (3.1) |  |
| Professional school degree | 364 (6.3) | 4 (0.4) |  |
| Doctoral degree | 403 (6.9) | 1 (0.1) |  |
| People living in home (mean (SD)) | 4.76 (1.64) | 4.97 (2.89) | 0.001 |
| Any siblings (yes, %) | 1905 (32.8) | 269 (26.0) | <0.001 |
| Hours/week spent at another household (mean (SD)) | 5.34 (19.45) | 5.45 (21.63) | 0.869 |
| Financial stress (0-7; mean (SD)) | 0.28 (0.85) | 1.32 (1.61) | <0.001 |
| Race (%) |  |  | <0.001 |
| Native American/Alaska Native | 17 (0.3) | 14 (1.4) |  |
| Asian | 126 (2.2) | 8 (0.8) |  |
| Black/African American | 495 (8.5) | 377 (36.5) |  |
| Pacific Islander | 8 (0.1) | 1 (0.1) |  |
| Other | 159 (2.7) | 74 (7.2) |  |
| White | 4263 (73.4) | 386 (37.3) |  |
| Mixed | 696 (12.0) | 141 (13.6) |  |
| Refuse to answer | 41 (0.7) | 33 (3.2) |  |
| Hispanic/Latino ethnicity (no, %) | 4776 (83.1) | 682 (67.3) | <0.001 |
| Parent marital status (%) |  |  | <0.001 |
| Married | 4621 (79.7) | 302 (29.6) |  |
| Widowed | 33 (0.6) | 22 (2.2) |  |
| Separated/divorced | 600 (10.4) | 232 (22.7) |  |
| Never married | 319 (5.5) | 369 (36.1) |  |
| Living with partner | 223 (3.8) | 96 (9.4) |  |
| Generational status (%) |  |  | <0.001 |
| Parent born outside U.S. | 708 (12.2) | 201 (19.5) |  |
| Grandparent born outside U.S. | 933 (16.1) | 90 (8.7) |  |
| Child born outside U.S. | 118 (2.0) | 32 (3.1) |  |
| Parents and grandparents born in U.S. | 4043 (69.7) | 709 (68.7) |  |
| School setting (%) |  |  | <0.001 |
| Not in school | 19 (0.3) | 6 (0.6) |  |
| Regular public school | 4836 (83.3) | 891 (86.2) |  |
| Regular private school | 346 (6.0) | 40 (3.9) |  |
| Charter school | 412 (7.1) | 79 (7.6) |  |
| Vocational/tech school | 2 (0.0) | 1 (0.1) |  |
| Cyber school | 7 (0.1) | 2 (0.2) |  |
| Home school | 112 (1.9) | 2 (0.2) |  |
| School for behavioral/emotional problems | 7 (0.1) | 3 (0.3) |  |
| Other | 63 (1.1) | 10 (1.0) |  |
| Youth-reported supportive school environment (6-24; mean (SD)) | 19.95 (2.63) | 19.96 (3.22) | 0.949 |
| Youth-reported school involvement (4-16; mean (SD)) | 13.11 (2.25) | 13.22 (2.44) | 0.162 |
| Youth-reported school disengagement (2-8; mean (SD)) | 3.66 (1.39) | 3.79 (1.57) | 0.006 |
| Census: % of people over age 25 with at least a high school diploma (mean (SD)) | 91.13 (8.76) | 81.30 (12.11) | <0.001 |
| Census: income disparity (mean (SD)) | 1.81 (1.17) | 3.13 (1.34) | <0.001 |
| Census: % of occupied units without complete plumbing (mean (SD)) | 0.28 (0.64) | 0.44 (0.83) | <0.001 |
| Census: % of families below the poverty level (mean (SD)) | 8.35 (8.68) | 20.93 (14.61) | <0.001 |
| Census: % of labor force aged >=16 y<br>unemployed (mean (SD)) | 7.69 (4.52) | 13.15 (7.49) | <0.001 |
| Census: uniform crime reports (mean (SD)) | 43774.47 (69634.30) | 43204.49 (57108.32) | 0.81 |
| Census: adult violent crime reports (mean (SD)) | 2660.87 (6271.58) | 2642.93 (5030.45) | 0.933 |
| Census: estimated lead risk<br>(1-10; mean (SD)) | 4.40 (2.98) | 6.77 (2.89) | <0.001 |
| Parent-reported neighborhood safety<br>(1-5; mean (SD)) | 4.05 (0.85) | 3.34 (1.11) | <0.001 |
| Parent self-reported aggressive behavior<br>(0-30; mean (SD)) | 3.14 (3.27) | 4.47 (4.58) | <0.001 |
| Parent self-reported intrusive behavior<br>(0-12; mean (SD)) | 1.01 (1.43) | 1.08 (1.43) | 0.198 |
| Parent self-reported withdrawn behavior<br>(0-18; mean (SD)) | 1.35 (1.85) | 2.46 (2.83) | <0.001 |
| Parent ethnic identification<br>(1-5; mean (SD)) | 2.71 (0.86) | 2.58 (0.94) | <0.001 |
| Youth-reported family conflict<br>(0-9; mean (SD)) | 1.93 (1.92) | 2.45 (2.04) | <0.001 |
| Youth-reported parental monitoring<br>(1-5; mean (SD)) | 4.43 (0.46) | 4.31 (0.59) | <0.001 |
| Youth-reported parental acceptance<br>(1-3; mean (SD)) | 2.80 (0.29) | 2.76 (0.33) | <0.001 |

Children’s scores on the three cognitive tests (Matrix reasoning, Flanker task, and Dimensional Change Card Sort task) were moderately correlated with each other, *r* = 0.23 – 0.43 in the whole sample, *r* = 0.25 – 0.39 for children living in poverty alone. We created summary cognitive test scores by summing children’s standardized scores on all three tests, as pre-registered. We first tested whether there was an association between income and cognitive test scores, using a linear mixed effects model with a random intercept for study site. For the purposes of comparison to prior studies, income was operationalized (for this analysis only) as a pseudo continuous variable, using the median income level in each income bracket. Results replicated prior studies (e.g., Duncan & Magnuson, 2012; Farah, 2018; Noble et al., 2015): on average, children whose families had higher incomes tended to perform better on cognitive tests, *B* = 0.008, SE = 0.0004, *p* < 0.001, *r* = 0.24, a moderate effect size, though it accounts for only 6% of the variance in children’s cognitive test scores. As shown in Figure 1, however, there was large individual variability in cognitive test scores within each income bracket. It is this individual variability we sought to explore further.

**Figure 1.**
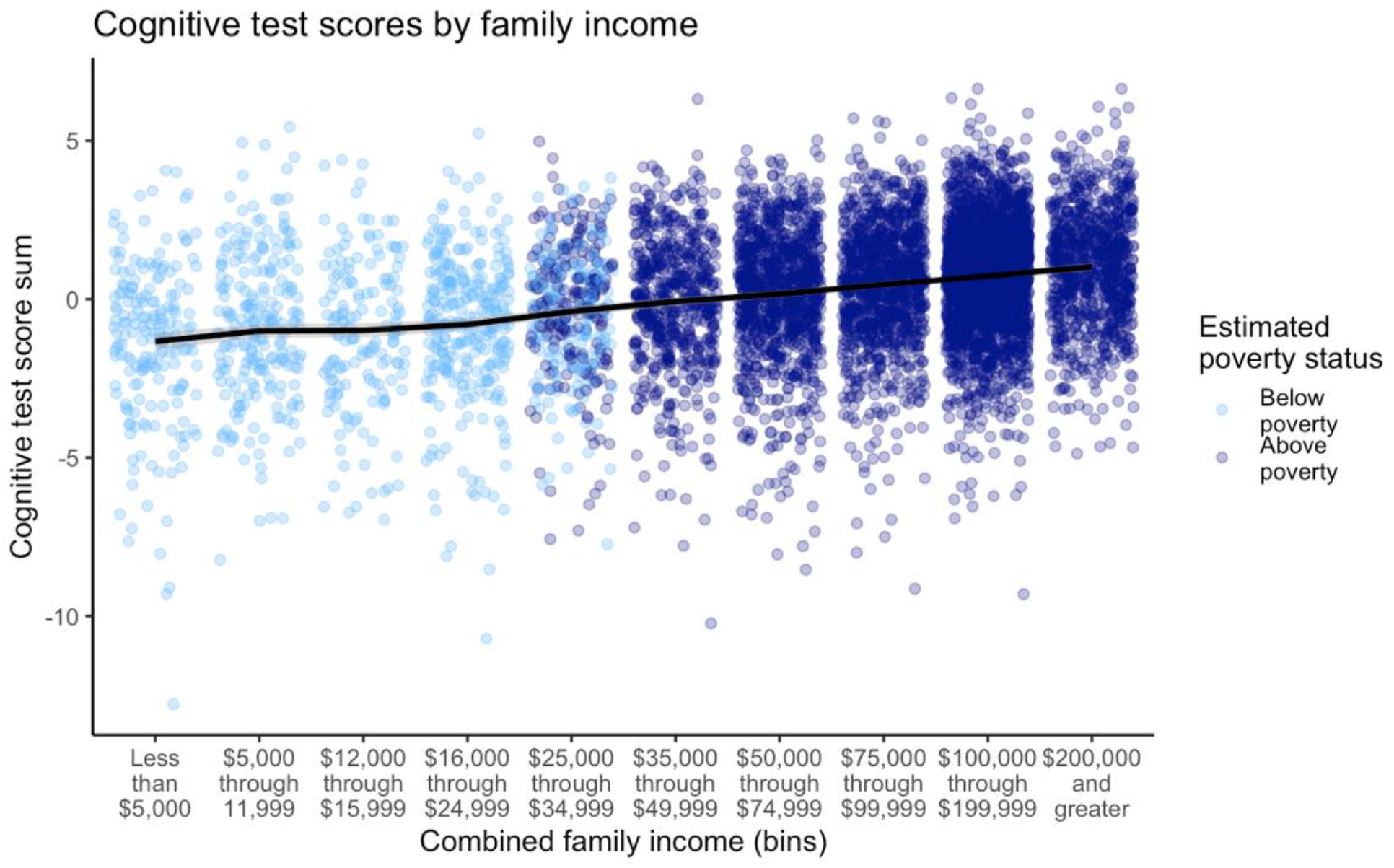
Illustration of the variability of cognitive test performance within every level of family income in the sample (N = 6839). Colors indicate whether children were classified as living in poverty, based on a combination of their family income and number of people in the home. Replicating prior studies, higher income is associated with higher cognitive test performance (R = 0.24); however, it is important to acknowledge this substantial variability within and overlap between children at each level of family income.

### LFPN-DMN connectivity

LFPN-DMN connectivity was defined as the average correlation of pairs of each ROI in LFPN with each ROI in DMN (each z-transformed; see Methods). Working from our pre-registered analysis plan (https://aspredicted.org/blind.php?x=3d7ry9), we tested the relation between LFPN-DMN connectivity and nonverbal cognitive test performance in our sample of children in poverty. We used linear mixed effects models to test the association between cognitive test performance and LFPN-DMN connectivity, controlling for children’s age and scanner head motion, with a random intercept for study site (see Methods). Contrary to previously published results, we did not find a negative association between LFPN-DMN connectivity and test performance. In fact, the estimated direction of the effect was positive, though this was not statistically significant, *B* = 2.11, SE = 1.12, *t* (1028) = 1.88; χ^2^ (1) = 3.52, *p* = 0.060. This numerically positive association was still observed when using a robust linear mixed effects model, which detects and accounts for outliers or other sources of contamination in the data that may affect model validity, *B* = 1.78, SE = 1.09, *t* = 1.64. Thus, this unexpected pattern was not driven by outliers. This effect was most pronounced for Matrix Reasoning and least evident for Flanker, but the estimate was positive for all three tests (see Supplement S2). It was also observed for the NIH Toolbox Fluid Cognition composite score (see Supplement S2).

Given this unexpected result, we next explored whether the expected association between LFPN-DMN connectivity and test performance was present in higher-income children in the larger dataset. To this end, we analyzed the 5,805 children from the same study sites who were likely to be living *above* the poverty line. Consistent with prior studies (Satterthwaite et al., 2013; Sherman et al., 2014; Whitfield-Gabrieli et al., 2020), these children showed a negative association between LFPN-DMN connectivity and cognitive test performance, *B* = −1.41, SE = 0.45, *t* (5794) = −3.14; χ^2^ (1) = 9.85, *p* = 0.002. A direct comparison between the samples confirmed that the association between LFPN-DMN connectivity and test performance differed as a function of whether or not children were living in poverty, χ^2^ (1) = 8.99, *p* = 0.003 (Figure 2). For children living above poverty, having higher LFPN-DMN connectivity appeared to be risk factor for low cognitive test performance, while for children living below poverty, this tended to be more protective. Several follow-up tests confirmed the reliability of this dissociation (see Supplement S4-S7). These included a bootstrapping procedure, permutation testing, and tests to ensure that results were not driven by differences in head motion, age, or the specific cognitive measures selected.

**Figure 2.**
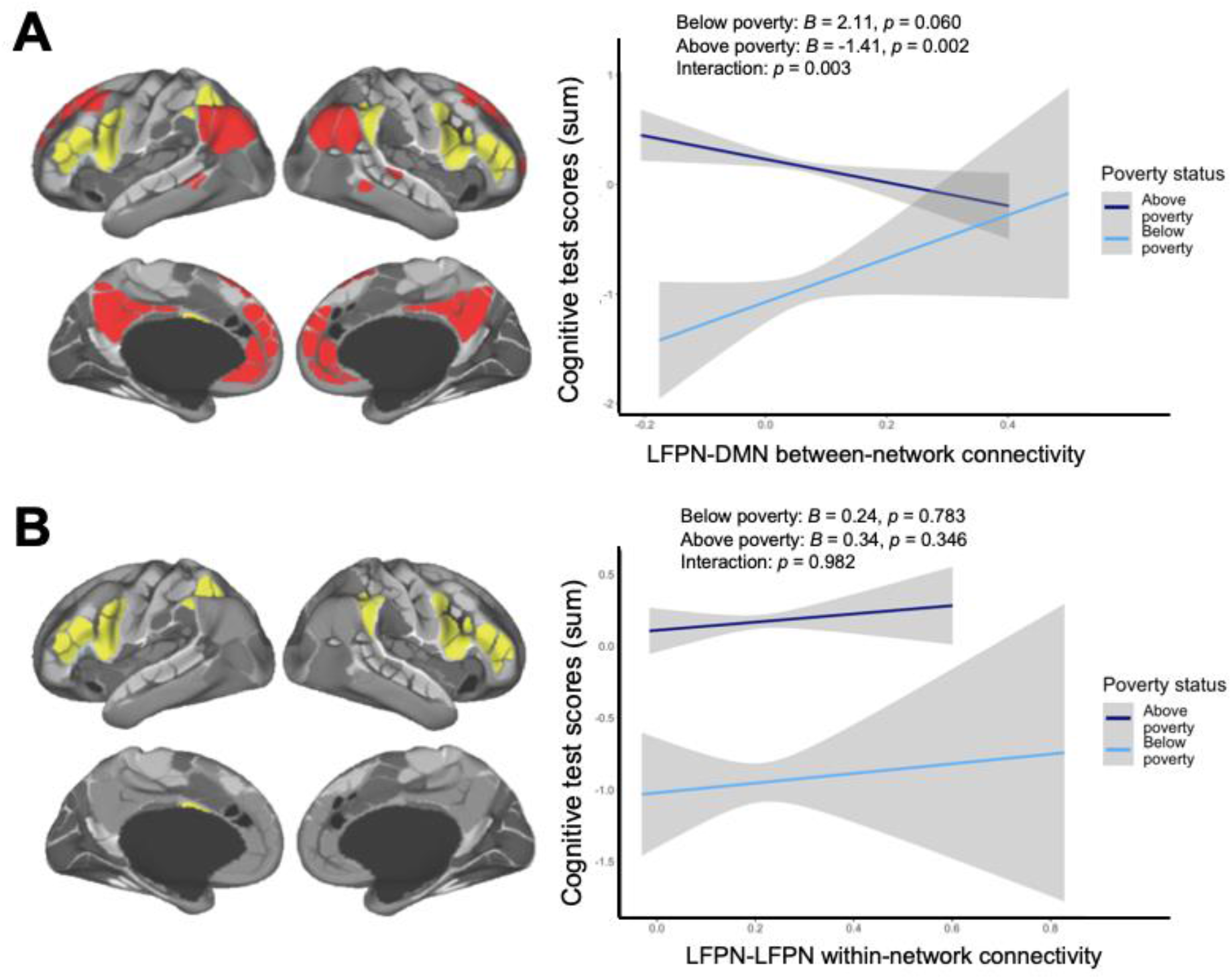
Relations between resting state network metrics and cognitive test score residuals, for children living above poverty (dark blue) and below poverty (light blue). Models include fixed effects for age and motion and a random effect for study site. 95% confidence intervals for a linear model calculated and displayed using the *geom_smooth* function in *ggplot*. Panel A: Children living above poverty show an expected, negative, relation between LFPN-DMN connectivity and test performance, *B* = −1.41, SE = 0.45; *p* = 0.002, while children living below poverty show the opposite pattern, *B* = 2.11, SE = 1.12; *p* = 0.060, interaction: *X*^2^ (1) = 8.99, *p* = 0.003. Panel B: Children across the sample show a non-significant positive relation between LFPN-LFPN within-network connectivity and test performance, above poverty: *B* = 0.34, SE = 0.36; *p* = 0.346; below poverty: *B* = 0.24, SE = 0.87; *p* = 0.783; interaction: *X*^2^ (1) = 0.0005, *p* = 0.982. Networks functionally defined using the Gordon parcellation scheme; on left, LFPN is shown in yellow and DMN shown in red, figures adapted from (Gordon et al., 2016).

### LFPN-LFPN connectivity

LFPN-LFPN connectivity was defined as the average correlation of each ROI pair within LFPN (each z-transformed; see Methods). Following our pre-registration, using linear mixed effects models, we next tested whether children in poverty would show the positive correlation between LFPN *within-network* connectivity and cognitive test performance that has previously been documented in higher-SES children. The relation between LFPN-LFPN connectivity and test scores was not significant for children in poverty, *B* = 0.24, SE = 0.87, *t* (1028) = 0.28; χ^2^ (1) = 0.08, *p* = 0.783, or for the higher income children in the larger study, *B* = 0.34, SE = 0.36, *t* (5797) = 0.94; χ^2^ (1) = 0.89, *p* = 0.346. Thus, strength of resting state functional connectivity within the LFPN network was not a predictor of cognitive performance in this large sample of 9 to 10-year-olds.

### Environmental variables

To further explore the dissociation observed for LFPN-DMN connectivity, we next asked whether features of children’s environments might explain why the brain-behavior link differed as a function of poverty status. Even among children living in poverty, different children are exposed to very different experiences in their homes, neighborhoods, and schools. Under what environmental constraints might it be optimal (with respect to cognitive test performance) for the LFPN to work more closely with the DMN? To answer this question, we considered 29 demographic variables chosen to reflect features of children’s home, school, and neighborhood environments (Table 2; see Appendix). To test whether any of these variables could explain the observed group interaction, we performed Ridge regression. Specifically, we used nested cross-validation to predict cognitive test performance from an interaction between LFPN-DMN connectivity and these demographic variables, in addition to main effects of each of these variables. Briefly, Ridge regression is a regularization technique that penalizes variables that do not contribute to model fit, thus giving more weight to the most important variables. This approach allows for the inclusion of many variables in a model while reducing the chances of overfitting, and deals with issues of multicollinearity. We pre-registered this second step of analyses prior to examining the data further (https://aspredicted.org/blind.php?x=tg4tg9), given the substantial analytic flexibility possible with such a large set of variables.

We trained our model in a training set of two-thirds (*N* = 670, after removing missing data) of the children in poverty, using 5-fold cross-validation. Next, we tested whether these demographic and neural model parameters could be used to predict cognitive test scores in the held-out test set: the remaining one-third (*N* = 329) of children in poverty. Indeed, we found that our model performed above chance (cross-validated R^2^_CV_ > 0; see Supplement S8), explaining 4% of the variance in children’s cognitive test scores in this held-out sample. While 4 percent is small, it is on par with the effect of family income on test scores across the full sample (6%). Additionally, it is a pure indicator, unlike the R^2^ of models that have been fit to the data themselves and are thus likely to be inflated. Most importantly, this prediction is based on a socioeconomically restricted sample of children: those with a total family income below $35,000 (below $25,000 for children in families of 4 or less).

As shown in Table 3, individual, home, neighborhood, and school variables helped to predict cognitive test scores among children living in poverty. Critically, we found that several characteristics of children’s experiences interacted with LFPN-DMN connectivity to predict these test scores. Specifically, variables related to school type, neighborhood safety, child’s race/ethnicity, and parents’ highest level of education contributed to model fit (see Table 3). To better understand these results, we plotted the effects for the factors showing significant interaction effects (Figure 3). Visualizing the interaction for neighborhood safety revealed that children living in safer neighborhoods showed a negative relation between LFPN-DMN connectivity and test performance, whereas those who lived in particularly dangerous neighborhoods showed a positive relation. With regard to schooling, the relation between LFPN-DMN connectivity was more positive for children attending public schools than those attending other types of schools (predominantly charter, *N* = 79, and private, *N* = 40). Thus, the brain-behavior relation for those children in poverty living in safer neighborhoods, or attending non-public schools, more closely resembled that of the higher-income sample. Similar results were obtained for levels of parental education and race, such that subsets of children whose parents were more highly educated and children who were white showed a more similar pattern of brain-behavior relations to children living above poverty.

**Figure 3.**
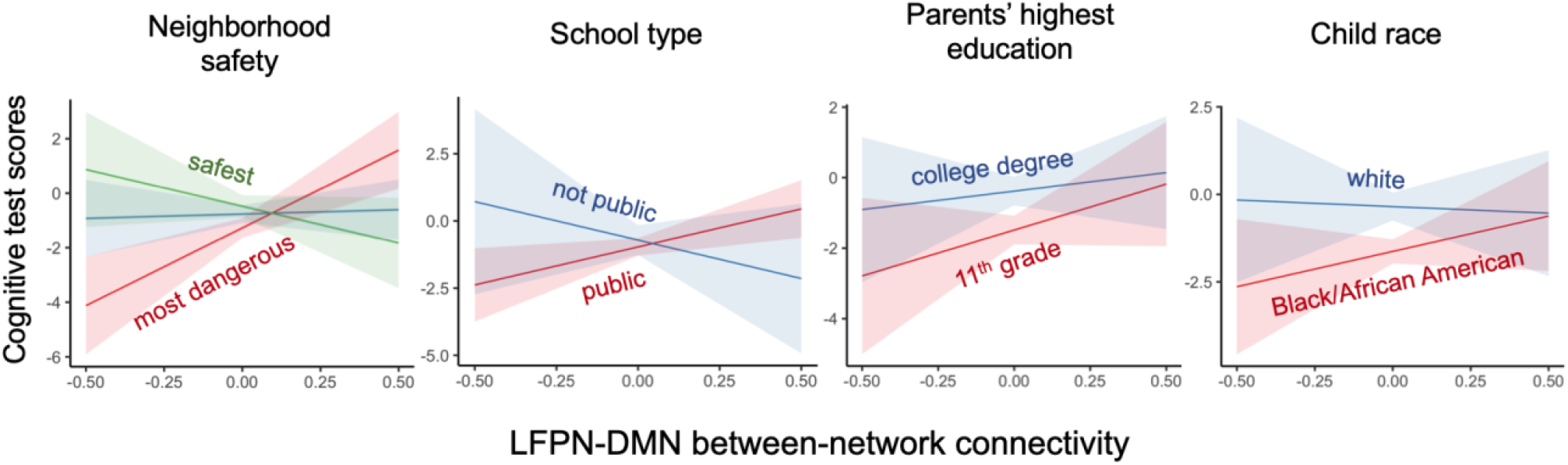
Interactions between demographic variables and LFPN-DMN connectivity in predicting cognitive test scores, for children below poverty. The majority of non-public schools were charter and private schools. In addition, only white and Black/African American race are displayed as these were the most represented in the current sample, though there were also suggestive interactive effects for children of mixed race and Hispanic ethnicity. 89% level confidence intervals for predicted effects calculated and displayed using the *sjPlot* package in R (Lüdecke, 2019).

**Table 3.**
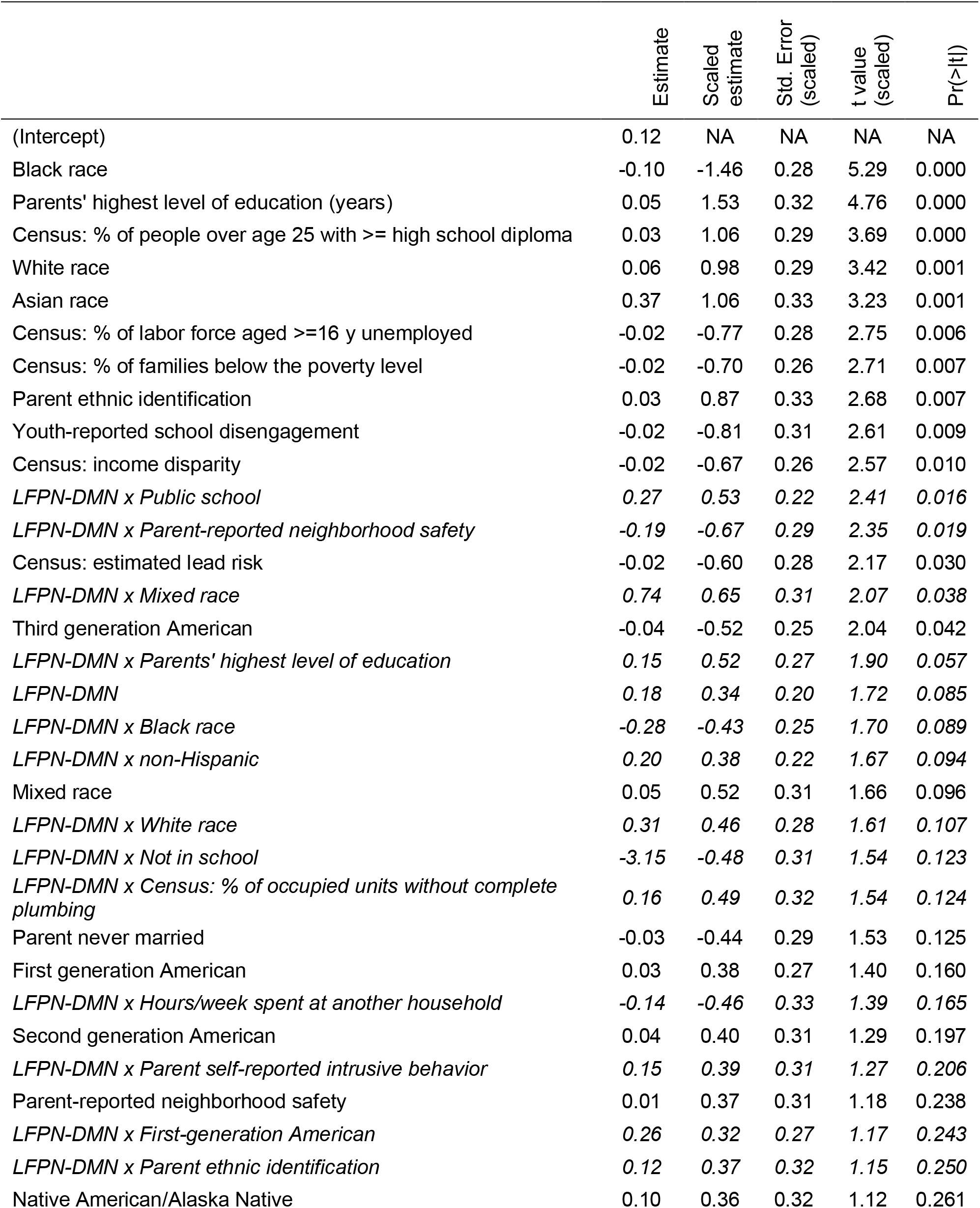
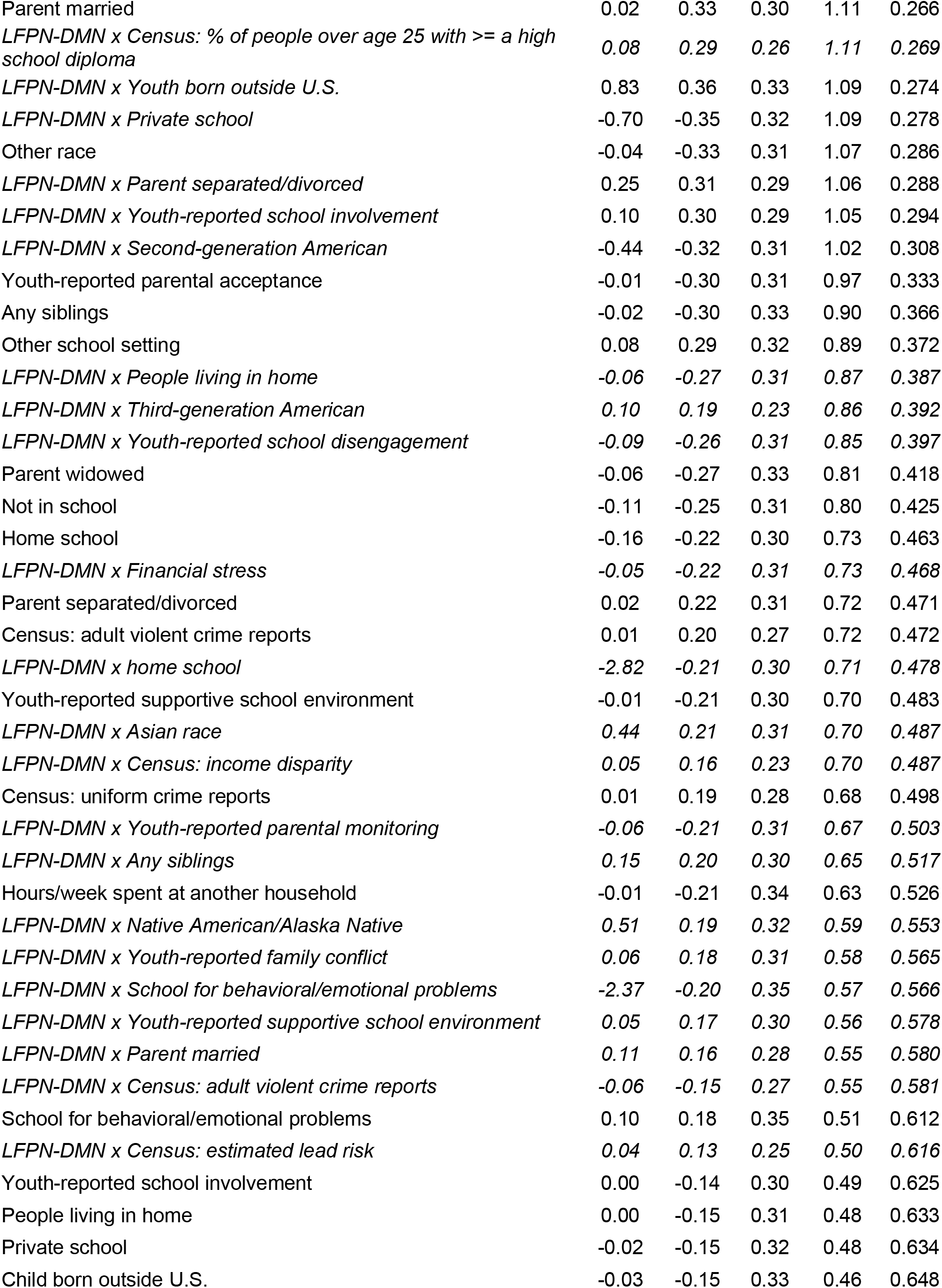
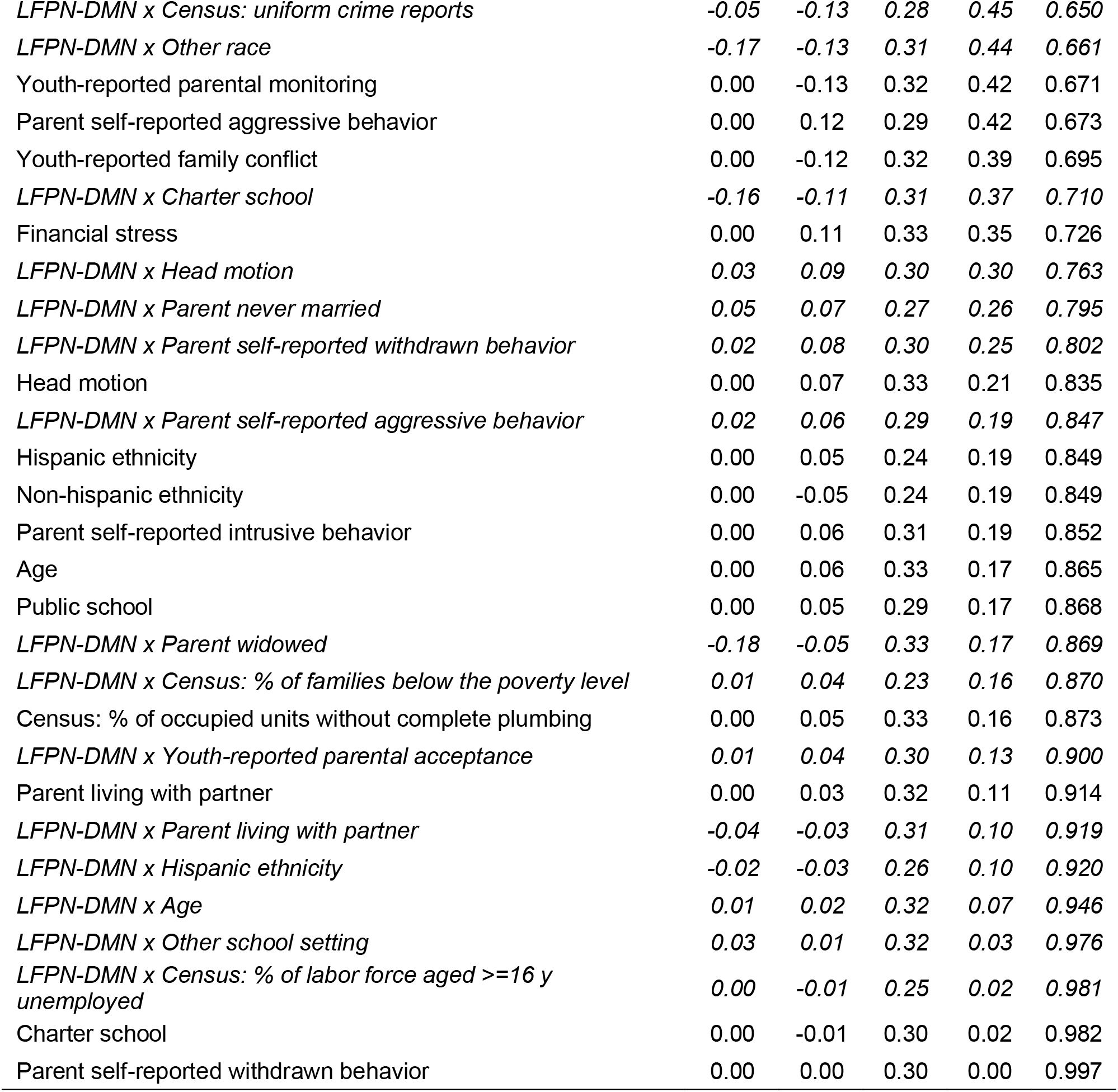
Estimated coefficients from Ridge regression predicting children’s cognitive test scores, when controlling for fixed effects of age and motion and random effects of study site, for all children below the poverty line. Interactions with and main effect of LFPN-DMN connectivity italicized.

Finally, we conducted a confirmatory factor analysis to test whether the demographic variables could be split into individual and home, neighborhood, and school factors based on our *a priori* categorization. This categorization did not meet our pre-registered criteria for a good model fit (our CFI, 0.11, was considerably lower than 0.9); as a result, we did not continue with this portion of the analysis. Thus, our data-driven approach provided insights that would have been missed by simply categorizing variables based on our prior assumptions about classes of life experiences.

### Exploratory network associations

Given the differential relation between network connectivity and test performance as a function of poverty status, we sought to ascertain whether this effect was specific to the LFPN-DMN, or whether there was a more general difference regarding connectivity between networks. Further, we sought to better understand the phenomenon at a conceptual level by assessing the plausibility of several accounts regarding what might constitute adaptive thought patterns for children contending with extremely challenging circumstances. Therefore, we ran several exploratory analyses involving two additional brain networks, selected for reasons discussed below. Due to the exploratory nature of these analyses, we focus on the general patterns of effects as potentially valuable for guiding future research.

The first additional network in which we tested for effects of poverty status was the cingulo-opercular network (CON), which is thought to play a role in coordinating the engagement of the LFPN and DMN networks (Menon & Uddin, 2010; Sridharan et al., 2008). Therefore, we sought to test for differential effects of coordination between the CON and these networks as a function of poverty. We found that weaker LFPN-CON connectivity was associated with better test performance for both groups, with little evidence of an interaction (Figure 4A). Thus, a dissociation between these networks appears to be generally adaptive at this age. By contrast, DMN-CON connectivity had no main effect on cognitive test performance, but it showed a possible interaction with poverty status (Figure 4B). Specifically, *weaker* DMN-CON connectivity was directionally associated with better test performance for children in poverty, while *stronger* DMN-CON connectivity appeared more adaptive for children above poverty. Thus, the cognitively adaptive pattern for children in poverty—at least, at this age (9-10)—is for DMN to be more tightly linked to LFPN and, perhaps, less tightly linked to CON. However, it seems unlikely that a DMN-CON interaction is the key driver of the LFPN-DMN interaction we have uncovered, as the latter effect was stronger. Nonetheless, further research in this population relating these three brain networks to a broader set of cognitive measures is warranted.

**Figure 4.**
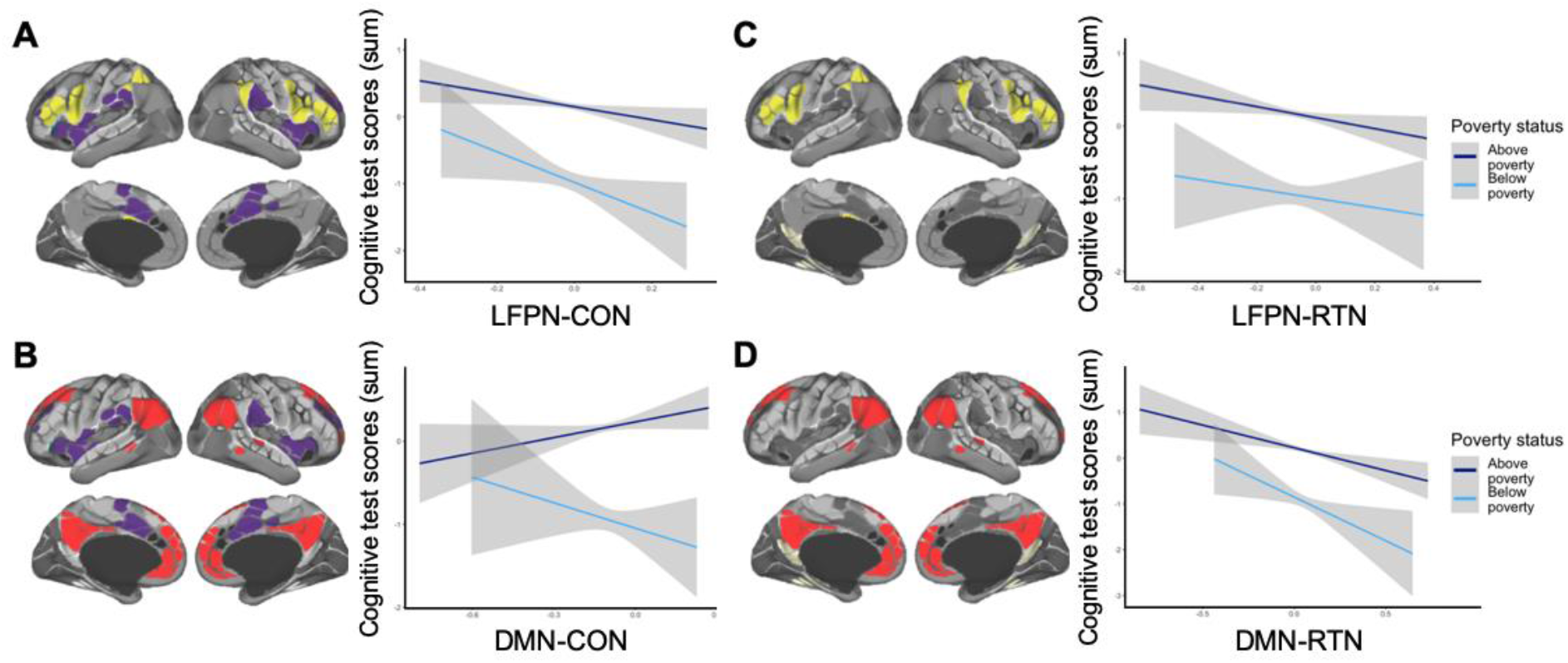
*Exploratory analyses with cingulo-opercular network (CON, panels A-B) and retrosplenial temporal network (RTN, panels C-D). **Panel A**: weaker LFPN-CON connectivity was associated with better test performance for both groups, with little evidence of an interaction (main effect: B = −1.14, SE = 0.45, t (6824) = −2.53;* X^2^ *(1) = 11.76, p = 0.001; interaction: B = −1.42, SE = 1.03, t (6824) = −1.37; X^2^ (1) = 1.87, p = 0.171). **Panel B**: DMN-CON connectivity was not consistently associated with test performance, though it was directionally positive for children above poverty and negative for children below poverty (main effect: B = 0.47, SE = 0.38, t (6823) = 1.24;* X^2^ *(1) = 0.27, p = 0.601; interaction: B = −1.66, SE = 0.88, t (6823) = −1.88;* X^2^ *(1) = 3.53, p = 0.060). **Panels C and D**: weaker LFPN-RTN connectivity and weaker DMN-RTN connectivity were both associated with better test performance, with little evidence of an interaction (**Panel C**: LFPN-RTN main effect: B = −0.90, SE = 0.36, t (6829) = - 2.54;* X^2^ *(1) = 7.13, p = 0.008; LFPN-RTN interaction: B = 0.23, SE = 0.84, t (6829) = 0.27;* X^2^ *(1) = 0.08, p = 0.784; **Panel D**: DMN-RTN main effect: B = −0.99, SE = 0.32, t (6826) = −3.14;* X^2^ *(1) = 16.24, p < 0.001; DMN-RTN interaction: B = −0.95, SE = 0.75, t (6826) = −1.27;* X^2^ *(1) = 1.61, p = 0.205). As in Figure 2, plots show r*elations between resting state network metrics and cognitive test score residuals, for children living above poverty (dark blue) and below poverty (light blue). Models include fixed effects for age and motion and a random effect for study site. 95% confidence intervals for a linear model calculated and displayed using the *geom_smooth* function in *ggplot*.

The other network we investigated was the retrosplenial temporal network (RTN), which is critical for long-term declarative memory (Ghetti & Bunge, 2012; Vincent et al., 2006). Regions in the RTN interact with the LFPN during performance of episodic memory tasks involving externally-presented stimuli (Badre & Wagner, 2007; Blumenfeld & Ranganath, 2007), but with the DMN during autobiographical memory retrieval (Andrews-Hanna et al., 2014; Buckner & Carroll, 2007; Kaboodvand et al., 2018) and at rest (Chai et al., 2014a), that is, during internally directed thought. We reasoned that if cognitively resilient children in poverty rely more on their autobiographical memory than do others when facing cognitive challenges, LFPN-RTN connectivity might be positively related to test performance in this sample. Contrary to this prediction, however, we found that *weaker* LFPN-RTN connectivity and DMN-RTN connectivity were associated with better test performance in both the below- and above-poverty samples (Figure 4C and 4D). Thus, these exploratory analyses involving the CON and RTN networks reveal specificity in the observed LFPN-DMN interaction effect.

## Discussion

Prior research in both adults and children suggests that, in order to perform well on cognitively demanding tasks, the LFPN must operate independently from the DMN (Chai et al., 2014b; Sherman et al., 2014; Whitfield-Gabrieli et al., 2020). Given that the LFPN and DMN have been linked to externally and internally focused attention, respectively, these findings are generally taken to suggest that it is optimal for individuals engaged in a cognitively demanding task involving externally presented stimuli to focus narrowly on the task at hand while inhibiting internally-directed or self-referential thoughts (Raichle et al., 2001; Simpson et al., 2001a, 2001b; D. H. Weissman et al., 2006). However, the majority of the research that led to this conclusion has been conducted with non-representative samples of individuals from higher-income backgrounds. Given the large heterogeneity of experiences and outcomes for children living in poverty, we focused on this relatively under-studied population.

In this study, we tested the relation between patterns of brain connectivity and nonverbal cognitive test performance for over 1,000 American children estimated to be living in poverty. Although children in poverty scored lower on average than their higher-income peers from the same study sites, there was large variability. Indeed, many of the children in poverty scored on par with children whose family incomes were considerably higher. In contrast to research with higher SES samples, we did not find that higher cognitive test scores were associated with stronger anti-correlations between the LFPN and DMN within this group; in fact, these children showed a non-significant positive relation between cognitive performance and functional connectivity between these networks. By contrast, for the children in the sample living above poverty, we replicated the negative relation observed in prior studies (e.g., Sherman et al., 2014). Thus, for children living above poverty, having higher LFPN-DMN connectivity could be a risk factor for lower cognitive test performance, while for children living below poverty, it could be protective.

Further confirming the reliability of this dissociation, both a bootstrapping analysis and permutation testing showed that models trained on the data from the children living above poverty did a poor job of predicting test performance for the children below poverty. It is important to note that the fact that we see statistically trending but numerically small group differences in overall LFPN-DMN functional connectivity, as well as no evidence of group differences in LFPN-LFPN connectivity. As such, the most salient difference between children below and above poverty in our analyses was not overall brain connectivity, but rather the relation between connectivity and cognitive performance.

This pattern of results is also in line with prior structural and task-based brain imaging studies showing interactions between SES and neural variables in relation to test performance (Leonard et al., 2019; Merz, Wiltshire, et al., 2019). For example, several studies have found SES differences in lateral prefrontal and parietal activation during cognitive tasks, core nodes of the LFPN (e.g., Finn et al., 2017; Sheridan et al., 2012). Together, these findings support the idea that which patterns of brain function are adaptive with respect to cognitive test performance depends on the environments that children must contend with.

One interpretation of this unexpected interaction is that the relation between LFPN-DMN connectivity and test performance depends in part on the demands of children’s daily experiences. It may be optimal under some circumstances to engage in thought patterns that more frequently co-activate the LFPN and DMN (e.g., Christoff et al., 2009; Fornito et al., 2012; Prado & Weissman, 2011). For example, while the DMN is generally thought to be suppressed during goal-directed tasks, it is in fact active during a variety of goal-directed tasks that require internal mentation, or projection outside of the here-and-now (Buckner & Carroll, 2007; Spreng, 2012). We return to this point later in the Discussion.

In contrast to our findings with LFPN-DMN connectivity, we found no significant association between within-network LFPN connectivity and test performance—either in the children living below or above poverty. These results were unexpected, given prior studies reporting that connectivity within the LFPN is positively related to cognitive test performance in both adults and children (Langeslag et al., 2013; Li & Tian, 2014; Sherman et al., 2014; Song et al., 2008). For example, Sherman and colleagues found that for 10-year-olds, higher IQ was correlated with higher connectivity between the dorsolateral prefrontal cortex and the posterior parietal cortex, two hub regions of the LFPN. One reason for the non-significant effect in our study may be that we examined connectivity within the LFPN as a whole, rather than looking at particular regions or subnetworks within LFPN. Thus, the entire network might not be developed enough by ages 9 to 10 to see this relation on a global scale.

To better characterize the positive relation between LFPN-DMN and test performance among the children living in poverty, we examined a number of demographic variables. While poverty status tends to be associated with a higher likelihood of particular experiences, such as racial or ethnic discrimination, more crowding in the home and financial strain, unsafe neighborhoods, and underfunded public schools, there is large variation in the experiences of children who live in poverty (DeJoseph et al., 2020). Moreover, experiences that are on average associated with worse cognitive outcomes (such as being deprived of caregiver support in early life) can, under some circumstances, produce *better* cognitive outcomes (Nweze et al., 2020), suggesting there may be different routes to achieving high cognitive performance in these cases. Thus, we predicted that differences in environmental influences *among* children in poverty would explain whether strong LFPN-DMN connectivity was adaptive or maladaptive for cognitive test performance.

Our analyses suggested that demographic variables could not be well fit to a pre-determined factor structure based on variables relating to the individual, home, neighborhood, and school; therefore, we took a data-driven approach to examine the effects of environmental variables. Because many of these variables are correlated with each other, we adopted an analytic approach—Ridge regression—that allows for collinearity. The results of this analysis suggested that, even within the population of children in poverty alone—children who are often conceptualized as a homogenous group—variation in their environments was predictive of their cognitive test performance. We note, however, that this was far from deterministic; a model trained on two-thirds of the children in poverty explained 4% of the variance in the held-out third, suggesting these variables accounted for a small amount of variance overall.

The most predictive variables in the model were main effects of children’s race/ethnicity, their parents’ highest level of education, and neighborhood-level characteristics such as the percent of people in their census tract who were unemployed, had not completed their high school degree by age 25, and were living in poverty. All of these variables reflect structural barriers that families may face, including access to resources and institutions, such as high-quality schools, jobs, and healthcare, stable housing in safe neighborhoods, and experiences of racism within these systems (Alexander, 2012; Chetty et al., 2018; Desmond & Kimbro, 2015; Kraus et al., 2019; Shedd, 2015). Thus, the strongest predictors of low-income children’s cognitive performance reflect structural constraints on children’s lives. However, our data also suggest that being raised by parents with strong ethnic identification may provide a psychological buffer against these and other threats, in line with other research (Cardoso & Thompson, 2010; Chen et al., 2015; Costigan et al., 2010; Simons et al., 2002; Varner et al., 2018).

Notably, we found—in addition to these main effects of demographic variables— several interactions between these variables and LFPN-DMN connectivity that predicted cognitive performance. While Ridge regression precludes us from drawing strong conclusions about the importance of specific variables, we highlight those that contributed significantly to model fit. For example, children in poverty who attended public schools, lived in subjectively more dangerous neighborhoods, and were Black (the next best represented racial group after white race in our sample below poverty) were more likely to show a positive relation between LFPN-DMN connectivity and test performance.

We considered several possible accounts of the current findings. One possibility is that in order to contend with structural barriers, children experiencing tremendous adversity in the form of poverty need to monitor their environments (vigilance), as well as their own behavior or performance (self-monitoring), to a greater degree than do other children. This hypothesis stems from research showing that individuals living in poverty are more likely to experience threat in the physical domain (safety; Friedson & Sharkey, 2015) or in the social domain (racism; Nuru-Jeter et al., 2009; Shedd, 2015); they are also likely to receive less direct feedback or instruction in crowded or underfunded public schools (Orfield & Lee, 2005; Reardon & Owens, 2014) and at home (McLoyd, 1998). Additionally or alternatively, children in poverty may benefit from thinking more about the past or the future—that is, drawing more on autobiographical memory and future-oriented thinking and planning (Buckner and Carroll, 2007)—or the type of productive mind-wandering that fuels creative insights (Christoff et al., 2009; Dixon et al., 2014; Seli et al., 2015). These hypotheses could be explored in the future by assessing whether children in poverty with stronger LFPN-DMN connectivity also show heightened self-monitoring, vigilance, autobiographical memory, and/or creative problem-solving.

Based on the available dataset, we explored the plausibility of these hypotheses by focusing on brain networks that have been associated with monitoring or declarative memory. Specifically, we explored associations of test performance with DMN/LFPN and (1) the cingulo-opercular (so-called “salience”) network (CON), to probe whether differences in monitoring and vigilance are likely to play a role; and (2) retrosplenial temporal network (RTN), to assess the plausibility of an account involving autobiographical memory or planning.

While relations with RTN and test performance did not distinguish the children above and below poverty, we observed a potential interaction between DMN-CON connectivity and poverty status in its association with test performance. Weaker DMN-CON appeared to be directionally associated with better test performance for children in poverty, and worse for children above poverty. Although it seems unlikely that this trend-level group interaction involving the CON is the key driver of the LFPN-DMN interaction we have uncovered, it does lend credence to the possibility that monitoring oneself and one’s social environment may be one mechanism through which children in poverty ultimately score highly on cognitive tests. It is also in line with work suggesting that CON plays a critical role in switching between LFPN and DMN activation (Sridharan et al., 2008), that connectivity between the three networks changes across age (Uddin et al., 2010), and that some social cognitive processes rely on all three networks (Schurz et al., 2020).

While our study benefited from the ABCD dataset’s rich objective measures of a child’s environment, there are other potential environmental and individual level variables that should be considered in future research (Bates et al., 2018; Merz, Wiltshire, et al., 2019; Pollak & Wolfe, 2020). Future research could also benefit from a more sensitive measure of poverty. Because the publicly available dataset did not specify which of the 19 study sites corresponded to which American city, as this was treated as protected information, we determined a cut-off for our poverty threshold based on cost-of-living across study sites. Because cities across the United States vary substantially in cost-of-living, we selected a stringent cutoff for the poverty line. Thus, there are almost certainly families in the above-poverty group that belong in the below-poverty group. If anything, therefore, the use of a more sensitive measure would likely magnify the group difference that we report. In addition, it is important to note that children’s performance on cognitive tests can fluctuate from day to day for a variety of reasons (Dirk & Schmiedek, 2016; Könen et al., 2015), including motivation (Somerville & Casey, 2010), which is a likely source of noise in our models.

Further, while we focused on three tests of non-verbal cognitive test performance, future studies should examine a broader range of cognitive systems, as these may be differentially affected by the environment (Rosen, Meltzoff, et al., 2019). For example, experiences of threat and deprivation have distinct effects on medial and lateral prefrontal cortex development, respectively (McLaughlin et al., 2019); these effects may be mediated in part by lower-level visual and attentional processes (Rosen, Amso, et al., 2019). Clearly, there is a need for research which investigates the precise mechanisms through which the environment affects specific neural and cognitive systems, particularly given that much of this environmental variation is still within a species-typical range of experiences (Humphreys & Salo, 2020). Overall, these results suggest that different patterns of brain activation for children living in poverty do not necessarily imply a deficit (Ellwood-Lowe et al., 2016). However, an important next step will be to follow these children longitudinally to see how LFPN-DMN connectivity and its relation with cognitive test performance changes across adolescence.

Another important area of research is to look beyond the canonical cognitive tasks used in the present study to identify assessments or testing contexts for which children living in poverty might be particularly adapted to excel (Frankenhuis et al., 2020). Doing so might reveal that some children who underperformed on the cognitive measures in the current study have strengths in other domains as a result of adaptation to their environments.

This study opens several questions about the neural underpinnings of these findings that should be further examined. Given individual variability in network topography (Seitzman et al., 2019), future studies should examine whether this variability contributes to our findings. In addition, LFPN and DMN are both summary network measures; there could be qualitative differences in node-to-node connectivity, or smaller interactions between sub-networks, that we are not capturing in the current study (Buckner & DiNicola, 2019; Dixon et al., 2018; Fornito et al., 2012; Lopez et al., 2020). Moreover, it would be helpful to look at children’s task-based activation and functional connectivity to examine whether children in poverty are more likely to activate DMN during neutral, externally driven cognitive tasks outside of their daily environments. Finally, given that these metrics only explain a small amount of variance, it is important to look at the contribution of other neural indices.

Given that the structures that govern success have been largely created around the needs of middle- and upper-middle class families, understanding the strengths of families in poverty—and how children may thrive in spite of these structural barriers—is critical. Altogether, these results highlight the substantial variability of experiences of children living in poverty, who are often conceptualized as a single, homogenous group and compared to higher-SES children. Moreover, they suggest that our field’s assumptions about generalizability of brain-behavior relations are not necessarily correct. Looking beyond convenience samples of children will ultimately lend more insight into the neural underpinnings of cognition, and may show that there is not a general guiding principle about what is optimal in the ways we have thus far assumed. Not only would this advance benefit developmental cognitive neuroscience as a field, but it may ultimately allow us to better serve disadvantaged youth.

## Methods

Analysis plans were pre-registered prior to data access (https://aspredicted.org/blind.php?x=3d7ry9, https://aspredicted.org/blind.php?x=tg4tg9) and analysis scripts are openly available on the Open Science Framework (https://osf.io/hs7cg/?view_only=d2acb721549d4f22b5eeea4ce51195c7). The original data are available with permissions on the NIMH Data Archive (https://nda.nih.gov/abcd). All deviations from the initial analysis plan are fully described in the Supplement S9.

### Participants

Participants were selected from the larger, ongoing Adolescent Brain Cognitive Development (ABCD) study, which was designed to recruit a cohort of children who closely represented the United States population (http://abcdstudy.org; see Garavan et al., 2018). This study was approved by the Institutional Review Board at each study site, with centralized IRB approval from the University of California, San Diego. Informed consent and assent was obtained from all parents and children, respectively. We planned to restrict our primary analyses to children who fell below the poverty line on the supplemental poverty measure, which takes into account regional differences in cost-of-living (Fox, 2017). For example, while the federal poverty level in 2018 was $25,465 for a family of four, the supplemental poverty level in Menlo Park, CA—one of the ABCD study sites—was estimated to be over $37,000 around the same time period. However, upon reviewing the data after our pre-registration, we found that study site in the ABCD data was de-identified for privacy reasons, and as a result we could not use study site-specific poverty cut-offs. Instead, we estimated each child’s poverty status based on their combined family income bracket, the number of people in their home, and the average supplemental poverty level for the study sites included in the sample.

Based on these factors, we considered children to be in poverty if they were part of a family of 4 with a total income of less than $25,000, or a family of 5 or more with a total income of less than $35,000. We made this determination by comparing children’s combined household income to the Supplemental Poverty Level for 2015-2017 averaged across study sites (Fox, 2017). We excluded children who did not provide information about family income and complete data on all three cognitive tests, and/or if their MRI data did not meet ABCD’s usability criteria (see below). In addition, due to a scanner error, we excluded post-hoc all children who were scanned on Philips scanners. This left us with 1034 children identified as likely to be living below poverty (6839 across the whole sample). Table 1 provides a breakdown of sample demographics.

### Cognitive test performance

Children’s performance was measured on three non-verbal cognitive tests. Specifically, children completed two tests from the NIH Toolbox (http://www.nihtoolbox.org): Flanker, a measure of inhibitory control (Eriksen & Eriksen, 1974), and Dimensional Change Card Sort (DCCS), a measure of shifting (Zelazo et al., 2013); and the Matrix Reasoning Task from the Wechsler Intelligence Test for Children-V (WISC-V), a measure of abstract reasoning (Wechsler, 2014). More details on each of these tests and their administration in the current study is described elsewhere (Luciana et al., 2018). These tests were chosen because they all tax higher-level cognitive skills while having relatively low verbal task demands. We created a composite measure of performance across these three domains by creating z-scores of the raw scores on each of these tests and summing them, as pre-registered; the tests were moderately correlated, 0.23 < *r* < 0.43, in the whole sample.

### MRI Scan Procedure

Scans were typically completed on the same day as the cognitive battery, but could also be completed at a second testing session. After completing motion compliance training in a simulated scanning environment, participants first completed a structural T1-weighted scan. Next, they completed three to four five-minute resting state scans, in which they were instructed to lay with their eyes open while viewing a crosshair on the screen. The first two resting state scans were completed immediately following the T1-weighted scan; children then completed two other structural scans, followed by one or two more resting state scans, depending on the protocol at each specific study site. All scans were collected on one of three 3T scanner platforms with an adult-size head coil. Structural and functional images underwent automated quality control procedures (including detecting excessive movement and poor signal-to-noise ratios) and visual inspection and rating (for structural scans) of images for artifacts or other irregularities (described in Hagler et al., 2019); participants were excluded if they did not meet quality control criteria, including at least 12.5 minutes of data with low head motion (framewise displacement < 0.2 mm).

### Scan parameters

Scan parameters were optimized to be compatible across scanner platforms, allowing for maximal comparability across the 19 study sites. All T1-weighted scans were collected in the axial position, with 1mm^3^ voxel resolution, 256 x 256 matrix, 8 degree flip angle, and 2x parallel imaging. Other scan parameters varied by scanner platform (Siemens: 176 slices, 256 x 256 FOV, 2500 ms TR, 2.88 ms TE, 1060 ms TI; Philips: 225 slices, 256 x 240 FOV, 6.31 ms TR, 2.9 ms TE, 1060 ms TI; GE: 208 slices, 256 x 256 FOV, 2500 ms TR, 2 ms TE, 1060 ms TI). All fMRI scans were collected in the axial position, with 2.4mm^3^ voxel resolution, 60 slices, 90 x 90 matrix, 216 x 216 FOV, 800ms TR, 30 ms TE, 52 degree flip angle, and 6 factor MultiBand Acceleration. Motion was monitored during scan acquisition using real-time procedures to adjust scanning procedures as necessary (see Casey et al., 2018); this prospective motion correction procedure significantly reduces scan artifacts due to head motion (Hagler et al., 2019).

### Resting state fMRI processing

Data processing was carried out using the ABCD pipeline and carried out by the ABCD Data Analysis and Informatics Core; more details are reported by Hagler et al. (2019). Briefly, T1-weighted images were corrected for gradient nonlinearity distortion and intensity inhomogeneity, and rigidly registered to a custom atlas. They were run through FreeSurfer’s automated brain segmentation to derive white matter, ventricle, and whole brain ROIs. Resting state images were first corrected for head motion, displacement estimated from field map scans, B0 distortions, and gradient nonlinearity distortions, and registered to the structural images using mutual information. Initial scan volumes were removed, and each voxel was normalized and demeaned. Signal from estimated motion time courses (including six motion parameters, their derivatives, and their squares), quadratic trends, and mean time courses of white matter, gray matter, and whole brain, plus first derivatives, were regressed out, and frames with greater than 0.2mm displacement were excluded. While the removal of whole brain signal (global signal reduction) is controversial in the context of interpreting anti-correlations (Chai et al., 2012; Murphy & Fox, 2017), we note that we are able to replicate prior studies showing that a more negative link between our networks of interest is related to test performance in our higher-income sample (see Results), lending credence to the inclusion of this step in the analysis pipeline for our purposes.

The data underwent temporal bandpass filtering (0.009 – 0.08 Hz). Next, standard ROI-based analyses were adapted to allow for analysis in surface space (Hagler et al., 2019). Specifically, time courses were projected onto FreeSurfer’s cortical surface, upon which 13 functionally-defined networks (Gordon et al., 2016) were mapped and time courses for FreeSurfer’s standard cortical and subcortical ROIs extracted (Desikan et al., 2006; Fischl et al., 2002). Correlations for each pair of ROIs both within and across each of the 13 networks were calculated. These were z-transformed and averaged to calculate within-network connectivity for each network (the average correlation of each ROI pair within the network) and between-network connectivity across all networks (the average correlation of pairs of each ROI in one network with each ROI in another network). Here, we examined only within-network connectivity for LFPN and between-network LFPN-DMN connectivity.

Altogether, the process for curbing potential contamination from head motion was three-fold. First there was real-time head motion monitoring and correction, as described above, and a thorough and systematic check of scan quality in collaboration with ABCD’s Data Analysis and Informatics Center. Second, signal from motion time courses was regressed out during preprocessing, and frames with greater than 0.2mm of framewise displacement were excluded from calculations altogether, as were time periods with less than five contiguous low-motion frames. Third, a final censoring procedure was employed to identify potential lingering effects of motion by excluding any frames with outliers in spatial variation across the brain (Hagler et al., 2019). In combination, these procedures reduce motion artifacts to the extent possible (Power et al., 2014).

### Analysis

Analyses were performed using R version 3.6.0 (R Core Team, 2017). We performed two separate linear mixed effects models using the *lme4* package (D. Bates et al., 2015) to test the relation between cognitive test scores and (1) LFPN-DMN connectivity, and (2) LFPN within-network connectivity. In our initial pre-registration, we did not consider the nested structure of the data or potential confounds. To determine whether to include these in our model in a data-driven fashion, we tested whether each of the following variables contributed significantly to model fit: (1) nesting within study site, (2) nesting within families, (3) child age, and (4) mean levels of motion in resting state scan. All except (2) contributed to model fit at a level of *p* < 0.01 and were thus retained in final models. We note that our reported results are similar when we perform simple linear regression with no covariates, exactly as pre-registered. In addition, results are similar when including all of the covariates in the ABCD study’s default LMM package (https://deap.nimhda.org/) – specifically, when adding fixed effects of race/ethnicity, sex, and parent marital status to the same model above. To determine the significance of our neural connectivity metrics, we tested whether these contributed to model fit. In all cases, we compared models without the inclusion of the variable of interest to models with this variable included, and calculated whether the variable of interest contributed significantly to model fit, using the *anova* function for likelihood ratio test model comparison.

In our second set of analyses, we sought to explore the unexpected results from our first set of analyses by asking whether certain environmental variables determine whether LFPN-DMN connectivity is positively or negatively associated with cognitive test performance across individuals. To do this, we gathered 31 environmental variables of interest, spanning home, neighborhood, and school contexts. Upon examining the data, we learned that three of these were not collected at the baseline visit and thus could not be included. Moreover, we made the decision to include ethnicity separate from race, as it was collected, to retain maximal information. The final 29 environmental variables are listed in Table 2. In preparation for our subsequent analyses, we mean-centered and standardized these variables in the larger dataset to allow for potential comparisons across the high- and low-income children. Levels of each factor variables were broken down into separate dummy-coded variables for inclusion in factor and ridge analyses. When data were missing, they were interpolated using the *mice* package in R (van Buuren & Groothuis-Oudshoorn, 2011).

We first performed a confirmatory factor analysis using the *lavaan* package in R (Rosseel, 2012) to see whether individual and home, neighborhood, and school variables can be separated into distinct factors. If this achieved adequate fit (significantly better fit than a single factor model and CFI>9), we planned to perform a linear mixed effects model to test the association of cognitive test performance with an interaction between LFPN-DMN connectivity and each factor score.

We next performed a ridge regression using the *glmnet* package in R (Friedman et al., 2010). This analysis technique penalizes variables in a model that have little predictive power, shrinking their coefficient closer to zero, thus allowing for the inclusion of many potential predictors while reducing model complexity. These models also include a bias term, reducing the chances of overfitting to peculiarities of the data, a common pitfall of ordinary least squares regression. Finally, ridge regression also deals well with multi-collinearity in independent variables; in contrast to alternatives such as Lasso, if two variables are highly correlated and both predictive of the dependent variable, coefficients of both will be weighted more heavily in ridge.

We fit ridge regressions predicting cognitive test score residuals, which partialled out the covariates included in our basic linear mixed effects models (random intercept for study site, fixed effects for age and motion), from an interaction between LFPN-DMN connectivity and each environmental variable of interest. This analysis used nested cross-validation. Specifically, we first split the data into a training (2/3) and testing (1/3) set. We created test score residuals in the training and testing sets separately to avoid data leakage (Scheinost et al., 2019), after rescaling the testing data by the training data. We then tuned parameters of the ridge regression on the training set using 5-fold cross-validation. Ultimately, we used the best-performing model to predict cognitive test scores in the held-out testing set and assessed model fit using R^2^ cross-validated. An R^2^_CV_ above 0 indicates that the model performed above chance; otherwise, it will be below 0. We evaluated the significance of specific variables in our model by plugging in the lambda parameter from the best-performing model to the linearRidge function in the *ridge* package in R (Cule & Moritz, 2019), on the whole sample of children in poverty.

### Robustness analyses

We did several additional analyses to test the robustness of our results. First, we repeated our primary analyses as robust linear mixed effects models, using the *robustlmm* package in R (Koller, 2016). These models detect outliers or other sources of contamination in the data that may affect model validity, and perform a de-weighting procedure based on the extent of contamination introduced. Next, we performed a bootstrapping procedure intended to probe how frequently the parameter estimate observed in the children in poverty alone would be expected to be observed in a larger population of children living above poverty (Supplement S4). We also performed a permutation procedure to examine the extent to which the model parameters from the higher-income children alone could explain the data in the children in poverty (Supplement S5). Finally, given that children living in poverty had significantly more motion than children living above poverty, we repeated our primary analyses with only those children who met an extremely stringent motion threshold of 0.2mm (Supplement S6).

Additional R packages used for data cleaning, analysis, and visualization include: *dplyr* (Wickham et al., 2019); *ggplot2* (Wickham, 2016); *car* (J. Fox & Weisberg, 2011); *corrplot* (Wei & Simko, 2017); *MuMIn* (Bartoń, 2019); *tidyr* (Wickham & Henry, 2019); *summarytools* (Comtois, 2019); *finalfit* (Harrison et al., 2019); *fastDummies* (Kaplan, 2019); *caret* (from Jed Wing et al., 2019); *scales* (Wickham, 2018); *foreign* (R Core Team, 2018); *MASS* (Venables & Ripley, 2002); *sjPlot* (Lüdecke, 2019); *tableone* (Yoshida, 2019); *gtools* (Warnes et al., 2018).

## Data availability

All raw and processed data used for these analyses are available with institutional permission on the NIMH Data Archive (https://nda.nih.gov/abcd).

## Code availability

All analysis scripts used for the current study are publicly available on the Open Science Framework (https://osf.io/hs7cg/?view_only=d2acb721549d4f22b5eeea4ce51195c7).

## Acknowledgements

This study would not be possible without the massive efforts of the large team of ABCD leaders and organizers, staff and data curators, and families and children who participated. Research reported in this publication also benefited from the ABCD Workshop on Brain Development and Mental Health, supported by the National Institute of Mental Health of the National Institutes of Health under Award Number R25MH120869. We are grateful to Mahesh Srinivasan for thoughtful comments on a previous draft of this manuscript, and to the members of the Building Blocks of Cognition Lab and the Language and Cognitive Development Lab for their feedback along the way. The content is solely the responsibility of the authors and does not necessarily represent the official views of the National Institutes of Health. MEL was supported by NSF GRFP DGE 1752814. SAB was supported by a Jacobs Foundation Advanced Career Research Fellowship.

