## Supplemental materials for "What is an adaptive pattern of brain network coupling for a child? It depends on their environment"

### 1. Scatterplots relating resting state metrics and cognitive test performance

For ease of viewing, Figure 2 in the main text displays trend lines of our primary models without the data points. Data points underlying Figure 2 are plotted in Supplementary Figure 1, below. This figure illustrates the extent of individual variability in the relation, and the sheer number of participants.

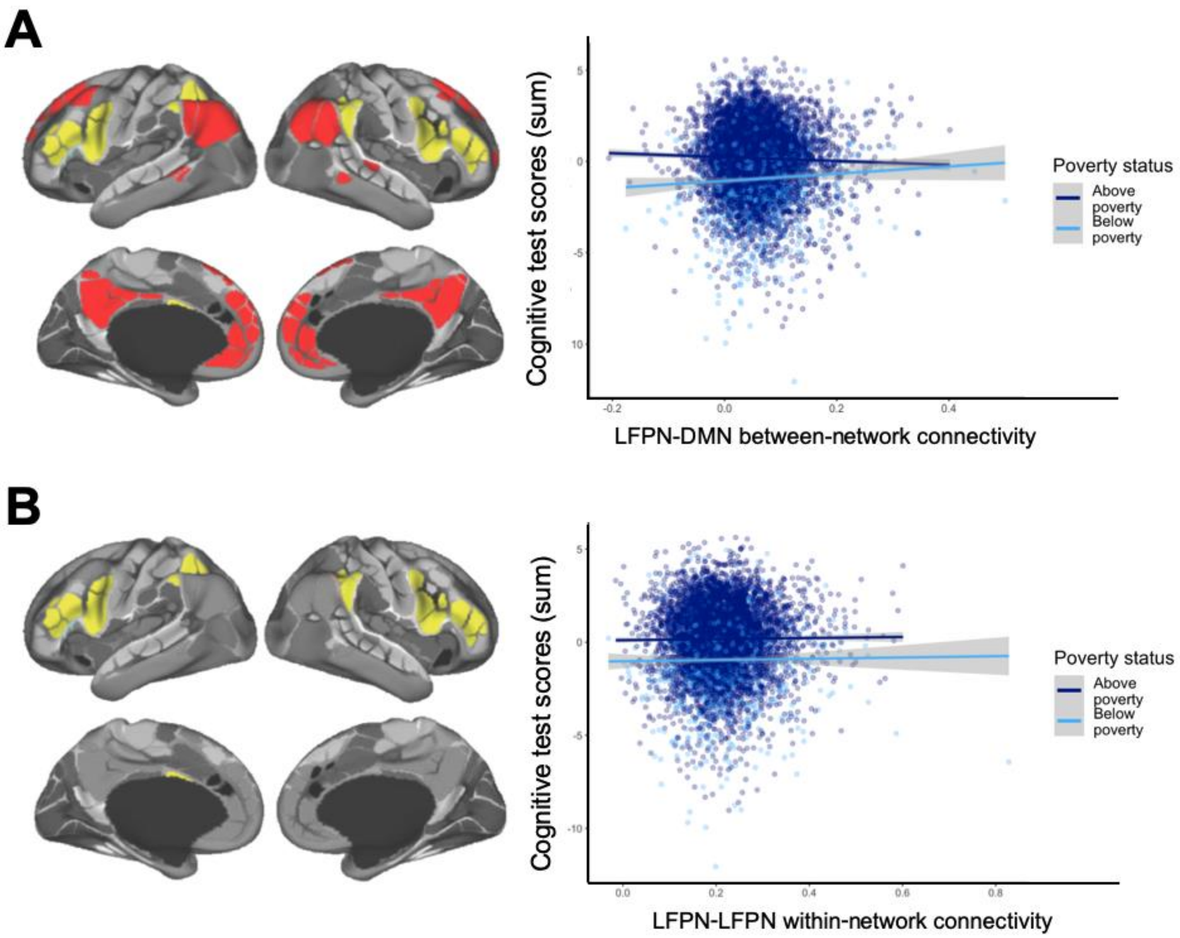

***Supplementary Figure 1****. Scatterplots with data points for relations between resting state network metrics and cognitive test score residuals, after accounting for fixed effects of age and motion and a random effect for study site, for children living above poverty (dark blue) and below poverty (light blue). Trend lines display 95% confidence intervals for a linear model using the* geom_smooth *function in* ggplot

### 2. Relations between LFPN-DMN connectivity and cognitive test performance, separated by test

Among the children in poverty, relations between LFPN-DMN connectivity and each cognitive test were in the positive direction, Matrix reasoning: *B* = 1.12, SE = 0.50, *t* (1028) = 2.25; $\chi^2$ (1) = 5.07, *p* = 0.024; Flanker: *B* = 0.18, SE = 0.56, *t* (1028) =

0.32; $\chi^2$ (1) = 0.1, $p$ = 0.751; Dimensional Change Card Sort (DCCS) task: $B$ = 0.81, SE = 0.51, $t$ (1028) = 1.6; $\chi^2$ (1) = 2.57, $p$ = 0.109.

In addition, mirroring our main results, this relation interacted with poverty status, reaching significance for reasoning, $\chi^2$ (1) = 6.76, $p$ = 0.009, and dimensional card sort, $\chi^2$ (1) = 6.44, $p$ = 0.011, but not for Flanker, $\chi^2$ (1) = 1.42, $p$ = 0.233.

We also repeated analyses using the NIH Toolbox Fluid Cognition composite, which includes two tests of working memory (Picture Sequence Memory Test, List Sorting Working Memory Test) and a test of processing speed (Pattern Comparison Processing Speed Test), in addition to Flanker and Dimensional Card Sort. (Note that Matrix Reasoning is not included in the fluid ability composite.)

Mirroring our primary results, relations between LFPN-DMN connectivity and the NIH Toolbox fluid ability composite were in the positive direction for children in poverty, $B$ = 9.19, SE = 4.95, $t$ (1020) = 1.86; $\chi^2$ (1) = 3.44, $p$ = 0.064, and negative for children above poverty, $B$ = -8.55, SE = 2.23, $t$ (5766) = -3.84; $\chi^2$ (1) = 14.7, $p$ < 0.001. Likewise, the relation between LFPN-DMN connectivity and the NIH Toolbox fluid ability composite interacted significantly with poverty status, $\chi^2$ (1) = 10.91, $p$ = 0.001, as shown in Supplementary Figure 2 below.

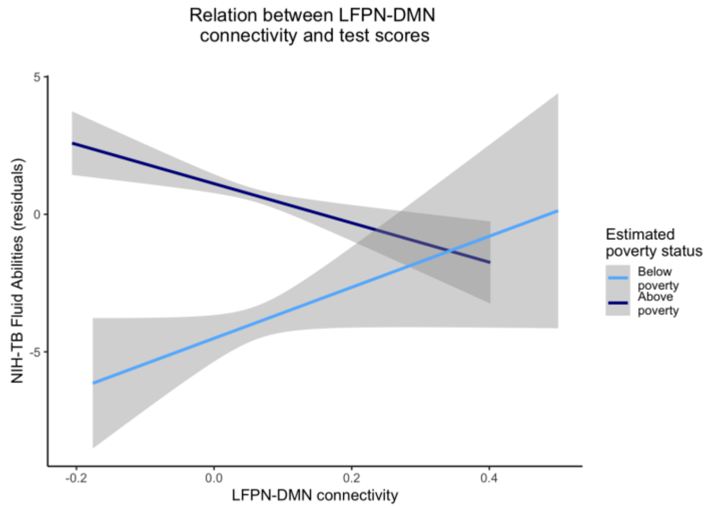

***Supplementary Figure 2.*** *Relations between LFPN-DMN connectivity and NIH-TB fluid abilities composite score residuals, for children living above poverty (dark blue) and below poverty (light blue). Models include fixed effects for age and motion and a random effect for study site.*

### 3. Relations between LFPN-LFPN connectivity and cognitive test performance, separated by test

Among the children in poverty, the direction of association between LFPN-LFPN connectivity and each cognitive test were inconsistent, matrix reasoning: $B$ = 0.69, SE = 0.38, $t$ (1028) = 1.81; $\chi^2$ (1) = 3.27, $p$ = 0.070; Flanker: $B$ = -0.44, SE = 0.42, $t$ (1028) = -

1.04; $\chi^2$ (1) = 1.06, $p$ = 0.303; dimensional card sort: $B$ = -0.11, SE = 0.39, $t$ (1028) = -0.28; $\chi^2$ (1) = 0.08, $p$ = 0.776.

There were no significant interactions between poverty status and LFPN-LFPN connectivity in predicting cognitive test scores, matrix reasoning: $\chi^2$ (1) = 2.36, $p$ = 0.125; Flanker: $\chi^2$ (1) = 0.78, $p$ = 0.376; dimensional card sort: $\chi^2$ (1) = 0.56, $p$ = 0.455.

### 4. Bootstrapped distribution of LFPN-DMN connectivity ~ test performance parameter estimates

Our first test was designed to probe how frequently the parameter estimate observed in the children in poverty would be expected to be observed in a larger sample of children living above poverty. In order to derive an estimate for observed parameter estimates in a population of higher-income children, we randomly sampled 500 data points from the children living above poverty, with replacement. For these 500 data points, we fit our primary linear mixed effects model to the data, predicting children's cognitive test scores, and calculated the average parameter estimate for LFPN-DMN connectivity. We repeated this process 999 times, generating a distribution of parameter estimates likely within the larger population of higher-income children from which our participants were drawn.

Next, we compared these bootstrapped parameter estimates to the parameter estimate observed for children in poverty in our sample. If the brain-behavior relation does not differ systematically as a function of poverty status—in other words, if the observed relation between LFPN-DMN connectivity and cognitive test scores for the children in poverty would be likely to be observed in a larger, population-level sample of children above poverty—the parameter estimate for children in poverty should fall within the 95% confidence interval of the bootstrapped parameter estimates.

Thus, we estimated the expected distribution of LFPN-DMN coefficients for the prediction of cognitive test scores among the higher-income children in the dataset. The results of this analysis confirmed that the observed estimate for children in poverty fell outside of the 95% CI, and was higher than 987 out of 999 bootstrapped samples, $p$ = 0.013.

Repeating this bootstrapping procedure for children living below poverty revealed a similar effect. Bootstrapped coefficients ranged from -2.87 to 7.66, with a mean of 2.26, 95% CI [2.16, 2.35]. The observed estimate for children above poverty fell outside of the 95% CI, and was lower than 990 out of 999 bootstrapped estimates, $p$ = 0.010. The bootstrapped distributions from the two samples are plotted side by side in Supplementary Figure 3.

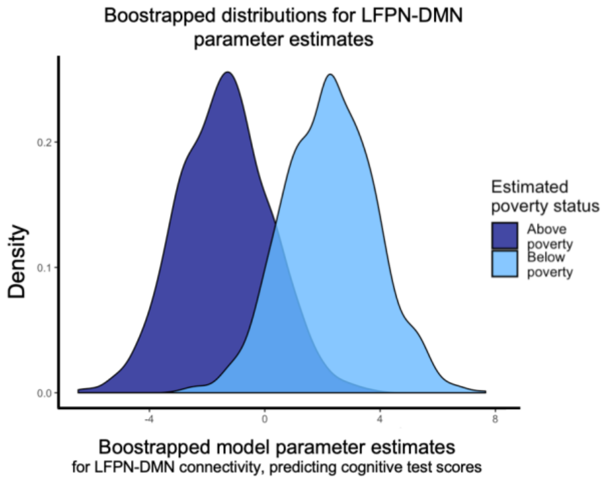

***Supplementary Figure 3.*** *Bootstrapped distributions for LFPN-DMN connectivity parameter estimates in the models predicting cognitive test performance, for children above (dark blue) and below (light blue) poverty. Estimates calculated from models run on 500 data points drawn with replacement from each sample separately, repeated 999 times. Bootstrapped coefficients for children above poverty ranged from -6.47 to 3.74, with a mean of -1.41, 95% CI [-1.50, -1.31], mirroring our observed parameter estimate for the higher-income group. The observed estimate for children in poverty fell outside of the 95% CI, and was higher than 987 out of 999 bootstrapped samples, p = 0.013.*

### 5. Permutation testing of LFPN-DMN connectivity ~ test performance parameter estimates

To further confirm the dissociation with LFPN-DMN connectivity and test performance for children living above or below poverty, we performed a permutation procedure. This procedure examined the extent to which the model parameters fit in the higher-income children alone could explain the data in the children in poverty.

Briefly, we used the model parameters generated from the higher-income children to predict test performance in the children in poverty, and calculated the mean difference (observed values minus predicted values). We next randomly permuted the labels of each group, such that assignment into the higher- versus lower-income group was now arbitrary. We repeated the process above, now fitting model parameters to our arbitrary “higher-income” group and using them to predict cognitive test performance in our arbitrary “lower-income” group. Again, we calculated the mean error between observed and predicted values. This permutation and prediction procedure was repeated 999 times, generating a distribution of mean differences when the distinction between the two groups was arbitrary. If the model parameters generated from our actual higher-income group could reasonably be applied to our actual lower-income group, we would expect that the mean error would fall within the 95% confidence interval of our distribution of permuted mean errors.

The results of this permutation procedure revealed that the model parameters fit on the children above poverty over-estimated the performance of children below poverty, on average (mean difference between observed and predicted test scores = -1.23). To contextualize whether this difference in prediction was larger than what would be expected by chance, we compared this to the distribution of 999 randomly group permuted labels, such that assignment into the higher- versus lower-income group was now arbitrary. The mean difference between the actual groups fell above all differences in the 999 permutations (range = -0.22 – 0.22), suggesting this difference is larger than would be expected by chance.

### 6. Relations between LFPN-DMN connectivity and cognitive test performance, for children with low thresholds of motion

Head motion, which is known to influence functional connectivity estimates (Power et al., 2015), differed significantly as a function of poverty status (see Table 1) and was correlated with cognitive test performance ($B$ = -1.91, SE = 0.16, $p$ < 0.001). Our reported analyses use a stringent motion exclusion criteria, in which participants were retained only if they had at least 12.5 minutes of data with low head motion (FD < 0.2 mm). Additionally, there was stringent motion correction in the analysis pipeline, as reported in the main text.

Still, we repeated these analyses with only those children who met a highly stringent motion criterion of less than or equal to 0.2 mm of average framewise displacement (N = 4444; 589 below poverty). Specifically, we fit a linear mixed effects model with site as a repeated measure, testing the relation between cognitive test scores and LFPN-DMN connectivity, controlling for age and head motion. In this subsample of participants who met our threshold for low motion (N = 4444; 589 below poverty), results were consistent, if not stronger. Specifically, for children living below poverty, the main effect of LFPN-DMN connectivity on test scores was positive and significant, $B$ = 4.92, SE = 1.92, $t$ (583) = 2.57; $\chi^2$ (1) = 6.61, $p$ = 0.010. Children living above poverty, in contrast, showed a negative main effect of LFPN-DMN connectivity, $B$ = -1.27, SE = 0.62, $t$ (3844) = -2.039; $\chi^2$ (1) = 4.15, $p$ = 0.041. The interaction between poverty status and LFPN-DMN connectivity was significant, $\chi^2$ (1) = 11.93, $p$ = 0.001 (consistent with the interaction effect in the full sample, $\chi^2$ (1) = 8.99, $p$ = 0.003). Thus, results were consistent—and seem to be even stronger—in this subsample of low-motion children.

### 7. Relations between LFPN-DMN connectivity and age

Given prior evidence that the LFPN and DMN become less correlated during childhood, we asked whether there was an effect of age in the current study. Indeed, even within this very restricted age range, LFPN-DMN connectivity was lower among older children across the entire sample, B = -0.0003, SE = 0.0001, t (6832) = -2.93 p = 0.003. This pattern was consistent both for children below poverty, B = -0.0004, SE = 0.0003, t (1032) = -1.66, p = 0.097, and those above, B = -0.0002, SE = 0.0001, t (5798) = -2.40, p = 0.016 (Supplementary Figure 4). We note that, while children in poverty had marginally higher LFPN-DMN connectivity overall (see Table 1), children living above poverty were approximately 17 days older than children living below poverty. The difference in connectivity between these groups was eliminated when accounting for this slight age difference (see Table 1).

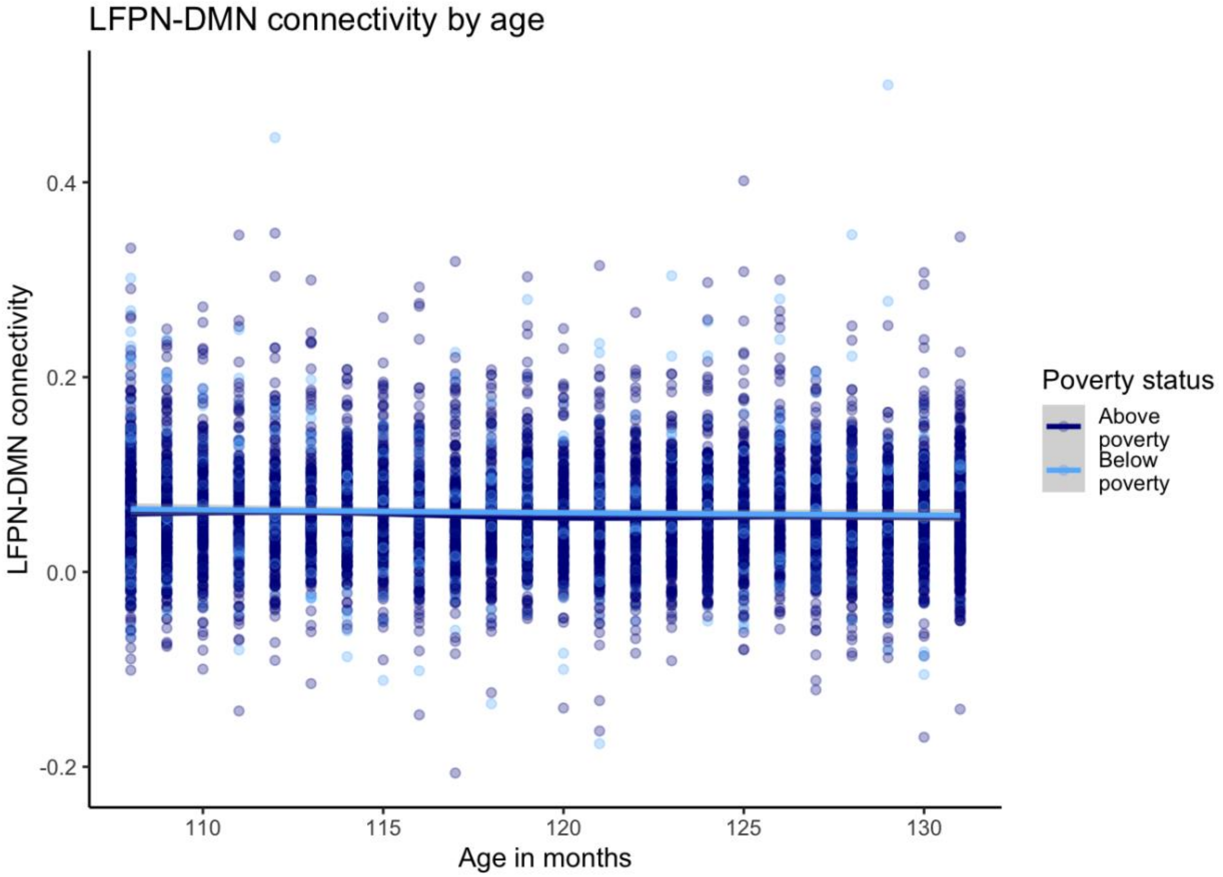

***Supplementary Figure 4***.
*Relations between LFPN-DMN connectivity and child age, for children estimated to be living above poverty (dark blue) and below poverty (light blue).*

### 8. Ridge regression confidence intervals

We randomly divided our test set and training set and found that it predicted approximately 4% ($R^2_{CV} = 0.037$) of the variance in test performance. However, one concern is that this split of the data happened to be particularly lucky, and unrepresentative of other possible splits. Therefore, we repeated this random split 1,000 times, training a new model on a randomly selected two-thirds of children and testing it in the remaining one-third of held out children. Thus, we were able to calculate a distribution of $R^2$ values. This repeated subsampling revealed that the model trained in two-thirds of the children in poverty predicted the held-out sample of children in poverty at above chance levels in more than 95% of iterations (in a cross-validation framework, this means $R^2 > 0$). The mean of this distribution was 0.023, 95% CI [0.021, 0.024]. Thus, while our split and model was on the high end of possible model fits, the model did consistently perform above chance.

### 9. Deviations from pre-registration

Both pre-registrations were written before knowing which data from the ABCD study would be available for analysis, which led to some necessary changes upon receipt of the data. In our first pre-registration, we planned to examine the relation between reasoning performance and specific node-to-node connectivity; however, only summary network measures had been released when we conducted our investigation. We also planned to look at test scores longitudinally, but found that only the first timepoint of cognitive assessments had been completed. Thus, we focused our analyses on one of our two primary planned questions. In addition, we planned to run simple linear regressions; these did not take into account the nested structure of the data, which we ultimately addressed in a data-driven fashion using linear mixed effects models, as described in the analysis section of the main text. The nested structure of the data also made our planned cross-validation approach less feasible, and we therefore did not cross-validate this first set of analyses. Finally, we planned to define our poverty threshold based on the Supplemental Poverty Threshold for each study site; however, due to privacy issues with de-identifying study site, we were only able to use a coarser threshold averaging across study sites. In our second pre-registration, we listed three environmental variables that were not collected at the baseline visit: self-reported discrimination, negative life events, and positive life events. Finally, we made the decision to include ethnicity separate from race, as it was collected, to retain maximal information. Otherwise, all analyses were performed as planned.

Several previously unspecified decisions were also made in the analysis process. First, we chose to use raw, rather than age-standardized, cognitive test scores. The rationale was that for using raw scores was that (1) the age range within our sample is relatively tight, (2) brain-behavior relations are of interest within the sample, not in relation to test norms based on a different sample of children, and (3) brain imaging data we are using aren't age normalized. Second, several factor levels within the environmental variables had a very low incidence in the whole sample, with less than 15 participants total (school setting: cyber school; school setting: vocational/tech school; race/ethnicity: Native Hawaiian); these were grouped into the "other" designation for the given factor, to allow for successful cross-validation when the sample was split further. Third, we made the decision to impute missing data from the environmental variables to preserve sample size.
